# Organization of Associating or Crosslinked Actin Filaments in Confinement

**DOI:** 10.1101/614354

**Authors:** Maral Adeli Koudehi, David M. Rutkowski, Dimitrios Vavylonis

## Abstract

A key factor of actin cytoskeleton organization in cells is the interplay between the dynamical properties of actin filaments and cell geometry, which restricts, confines and directs their orientation. Crosslinking interactions among actin filaments, together with geometrical cues and regulatory proteins can give rise to contractile rings in dividing cells and actin rings in neurons. Motivated by recent in vitro experiments, in this work we performed computer simulations to study basic aspects of the interplay between confinement and attractive interactions between actin filaments. We used a spring-bead model and Brownian dynamics to simulate semiflexible actin filaments that polymerize in a confining sphere with a rate proportional to the monomer concentration. We model crosslinking, or attraction through the depletion interaction, implicitly as an attractive short-range potential between filament beads. In confining geometries smaller than the persistence length of actin filaments, we show rings can form by curving of filaments of length comparable to, or longer than the confinement diameter. Rings form for optimal ranges of attractive interactions that exist in between open bundles, irregular loops, aggregated and unbundled morphologies. The probability of ring formation is promoted by attraction to the confining sphere boundary and decreases for large radii and initial monomer concentrations, in agreement with prior experimental data. The model reproduces ring formation along the flat axis of oblate ellipsoids.

## 1 Introduction

The cellular activity of cytoskeletal filaments is regulated by a combination of biochemical and physical cues, which can give rise to cytoskeletal networks with varied functional organization, including dendritic protrusive networks, the mitotic spindle, and contractile actomyosin bundles. Cell geometry and confinement plays a major role, since actin filaments polymerize near cell membranes and form bundles that reach lengths of order 10 *µm*, comparable to the size of cellular organelles and the size of plant, animal and yeast cells. For example, crosslinking interactions among actin filaments, together with geometrical cues and regulatory factors can give rise to contractile rings in dividing cells [Pollard and Wu, 2010, Lee et al., 2012, Balasubramanian et al., 2012] and actin rings in neurons [Leite and Sousa, 2016]. The length of actin filaments in the fission yeast contractile ring (~ 1 *µm*) is comparable to the cellular diameter (4 *µm*) and both are shorter than the persistence length of bare actin filament (~ 10 − 17 *µm*).

A large number of experimental and theoretical studies of cytoskeletal filaments in purified systems, with controlled interactions with boundaries and/or confining environment have aimed to uncover basic principles of cytoskeletal organization [Honda et al., 1999, Limozin and Sackmann, 2002, Hase and Yoshikawa, 2006, Claessens et al., 2006b, Claessens et al., 2006a, Reymann et al., 2012, Deshpande and Pfohl, 2015, Tsai and Koenderink, 2015, Miyazaki et al., 2015]. Actin rings have been reconstituted in vitro by polymerization of actin filaments in cell-sized droplets or liposomes, when attraction between filaments was promoted through the Asakura-Oosawa depletion force or through the addition of crosslinker proteins [Limozin and Sackmann, 2002, Claessens et al., 2006a, Tsai and Koenderink, 2015, Miyazaki et al., 2015]. Further addition of myosin motors promoted ring formation and triggered ring contraction [Miyazaki et al., 2015].

These recent experimental works motivated us to perform computer simulations in order to better understand the conditions under which associating or crosslinked semi-flexible filaments establish rings in a confining environment. We focus on confinement to compartments that are not deformable and on the effects of passive crosslinking (for the purposes of this work do not consider the effects of motor pulling). Despite many theoretical works examining either the properties of single filaments in confining space or else associating/crosslinking filaments in a bulk, the combination of association/crosslinking plus confinement has not been examined in detail. While Miyazaki et al. [Miyazaki et al., 2015] identified the ratio of persistence length to confinement radius as the main regulating factor for ring formation, we show that interaction strength (between filaments and surface) and filament length are also important. Our study focuses on actin filaments, though many of our conclusions should apply to other semiflexible biopolymers such as microtubules, DNA, and RNA whose contour lengths can also be comparable to the size of their cellular compartment.

Before describing our method and results, we provide an overview of related prior works that show different tendencies for loop/ring/network structure formation under different conditions. Isolated single filaments in a bulk can form loops and other shapes by folding back on themselves. The equilibrium and kinetically-trapped structures of single collapsed semi-flexible polymers has been studied extensively theoretically, see [Wu et al., 2018] and references within. Two major condensed structures, rings and racquets, were observed using fluorescence microscopy for polymerizing actin in presence of divalent ions or depletant [Tang et al., 2001, Lau et al., 2009]. Numerical simulations showed the folding and unfolding process of a single self attractive filament can be described by a kinetic theory of toroid nucleation and growth [Yoshinaga, 2008].

Theoretical and experimental studies have shown how confinement of long semiflexible filaments causes filament bending and looping. Gârlea [Gârlea, 2015] studied theoretically the shapes of single cytoskeletal chains confined in spheres and ellipsoids of size smaller or comparable to the filament persistence length. The authors described a change in filament distribution with increasing filament length: short filaments avoid the boundary while longer filaments bend and adopt the shape of the confinement to minimize bending energy. Non-interacting filaments that are long compared to the confinement radius loop around the boundary, as described by analytical theory and simulations [Lagomarsino et al., 2007, Morrison and Thirumalai, 2009, Fošnarič et al., 2013, Vetter et al., 2014] Additional regimes such as semi-dilute and liquid crystalline regimes arise when considering long semi-flexible filaments in confining spaces with dimensions larger than the persistence length [Sakaue, 2007].

Much more work has been performed on interacting filaments in the bulk, which, by contrast to the single or confined filaments studies mentioned in the two preceding paragraphs, tend to form networks rather than loops or ring structures. The architecture of a mixture of actin and crosslinkers can be broadly classified as a crosslinked network of single filaments or a bundle network [Lieleg et al., 2010]. The mechanical properties of the network depend on mesh size, entanglement length, crosslinker distance, and bundle stiffness [Lieleg et al., 2010]. Small crosslinking proteins such as scruin [Shin et al., 2004] or fascin [Lieleg et al., 2007] favor bundle formation while larger crosslinkers such as *α*-actinin [Tempel et al., 1996, Lieleg et al., 2009] or filamin [Schmoller et al., 2009] have more complex structures, including composite networks of bundles and single filaments and bundle asters. Dense networks of actin filaments in bulk also show liquid crystalline behaviour [Kas et al., 1996, Zhang et al., 2017].

Depletion agents such as PEG create bundles, the thickness of which increases linearly with the depletion agent concentration [Tharmann et al., 2006]. It has been demonstrated that the structure of crosslinked actin networks may reflect dynamically arrested states, the structure of which depends on the kinetics of polymerization and bundling by diffusion [Falzone et al., 2012, Foffano et al., 2016].

Many theoretical and computational studies of bulk networks have explored the viscoelastic properties of crosslinked actin gels [Broedersz and MacKintosh, 2014, Kim et al., 2009], phase transitions from homogeneous-isotropic to lamellar network as a function of number of binding proteins and binding angle [Muller et al., 2015], structural transition kinetics [Picu and Sengab, 2018], and regimes in bundle mechanics as function of filament length and crosslink density [Bathe et al., 2008].

To explore the combined effect of confinement and filament association, we performed coarse-grained simulations by representing individual filaments as beads connected by springs (Fig. 1A). This coarse-grained approach allows us to simulate realistic confining diameters of a few *µm* and actin filament concentration in the *µM* range though not reaching concentrations where excluded volume interactions would cause nematic ordering (dense confined semi-flexible polymers were examined by Milchev et al. [Milchev et al., 2017]). We also consider cases where the confining size is smaller or comparable to the filament persistence length, as is the case in many cellular systems as well as prior experiments in vitro.

**Figure 1:**
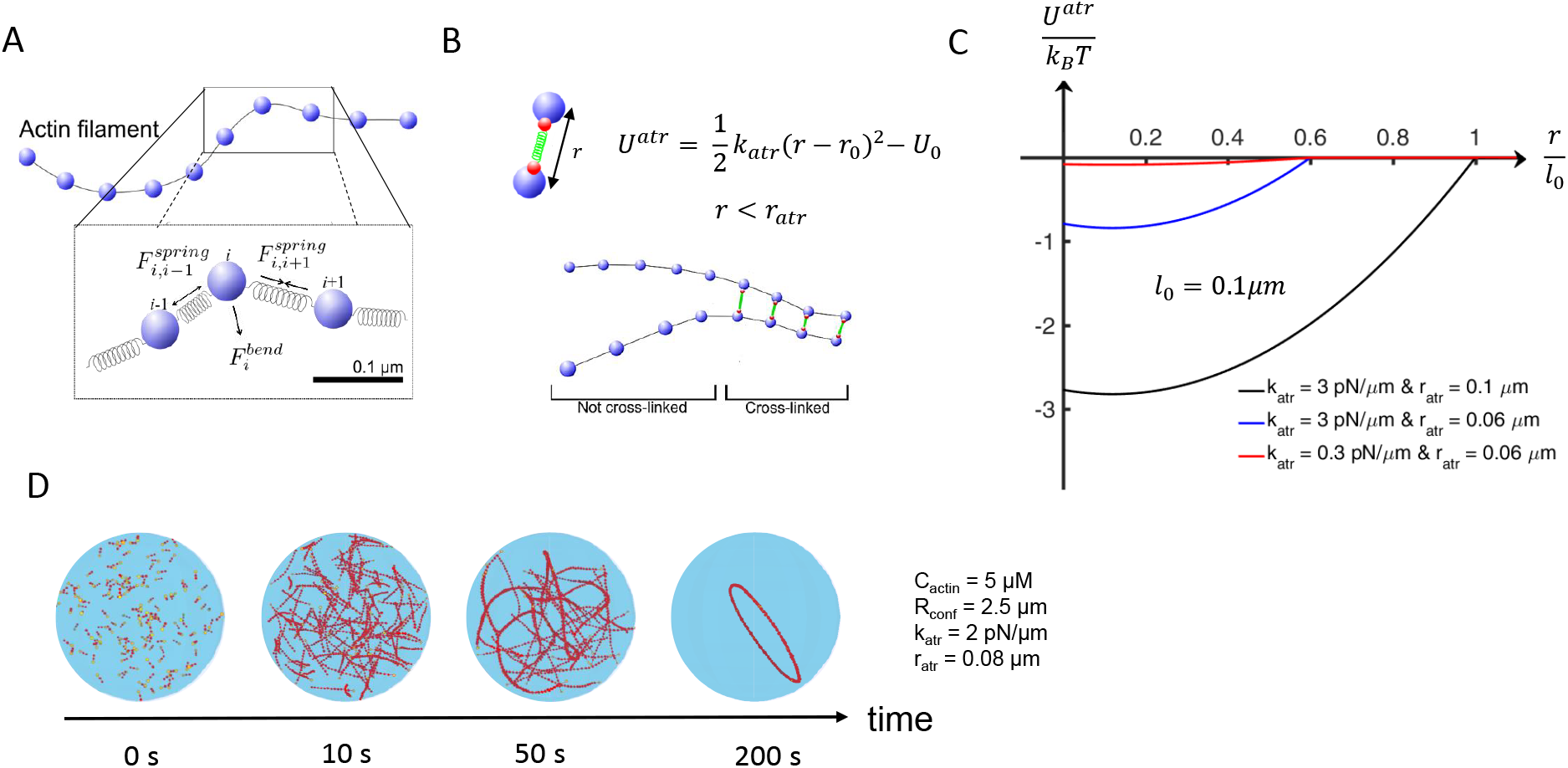
Simulation of attractive actin network in confinement. (A) A semi-flexible actin filament is described by a bead-spring model with spring and bending forces. The beads are separated by equilibrium distance *l*_0_ = 0.1 *µm*. (B) Attractive or crosslink interactions are represented by a force between two non-neighboring beads at distance *r* (when *r < r_atr_*) with spring constant *k*_*atr*_ and equilibrium length *r*_0_. (C) Plot of attractive potential for the indicated values of *l*_0_, *k*_*atr*_, and *r*_*atr*_ for *r*_0_ = 12 *nm*. (D) Example of actin ring formation in a confining droplet at indicated parameter values. In this simulation 140 filaments start to polymerize at *t* = 0 at a rate corresponding to the calculated concentration of remaining actin monomers in the bulk, reaching a final length of 3.8 *µm* after approximately 100 *s*. In snapshots of this and following figures, yellow beads are at the barbed ends.

In our model, filament association through the Asakura-Oosawa force or protein crosslinking is represented by a short range interaction potential between the actin filament beads. This interaction is described by two parameters controlling the width and depth of the potential (Fig. 1B,C). While this interaction does not accurately describe either depletion effects (that would be applied uniformly along the filament rather than at the discrete locations) or crosslinking (that depends on the helical pitch of the actin filament, for example), it has the advantage of providing a simple mechanism to tune the binding strength as well as filament sliding. We find a range of equilibrium and kinetically trapped states of confined filament organization that depend on concentration, length, confinement size, interaction strength among filaments and between filaments and boundary. This is thus a first step towards future work that could examine further important details such as the influence of specific crosslinker properties, including concentration, size, binding and unbinding kinetics.

## 2 Model

### 2.1 Filament representation

Following previous works [Tang et al., 2014, Bidone et al., 2014], each filament bead represents a segment of length *l*_0_ of the helical actin filament that includes 37 subunits. Bending energy and stochastic forces are included and these forces together with the boundary and crosslinking forces govern motion, using Brownian dynamics to update the 3D positions **r**_*i*_ of the *i*^th^ bead in time:

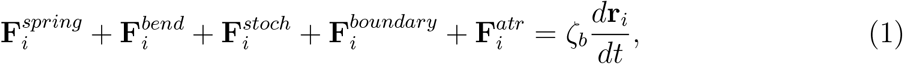

where *ζ_b_* is an effective drag coefficient. We use a drag coefficient close that to the viscosity of the cytoplasm (350 times higher than water [Tang et al., 2014]) and close to the viscosity of methlycellulose solutions in [Miyazaki et al., 2015], estimated to be 100 times higher than water (based on the stated concentration and Sigma-Aldrich specifications). For this viscosity we use an integration time step *dt* = 1.5 × 10^*−*4^*s*. We checked that the simulations were stable up to *dt* = 2 × 10^*−*4^*s* but became unstable for *dt* = 3 × 10^*−*4^*s*. Thus the selected time step was below the stability threshold while smaller values did not lead to changes in the results. Simulations of less viscous solutions are beyond the reach of our current method as they would require a significantly smaller *dt*.

The spring, bending and stochastic forces are as follows (Fig. 1A) [Tang et al., 2014, Pasquali et al., 2001]:

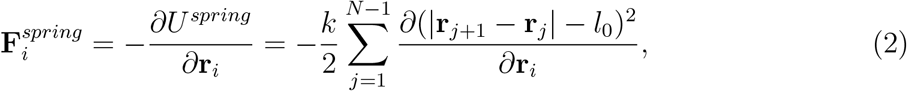

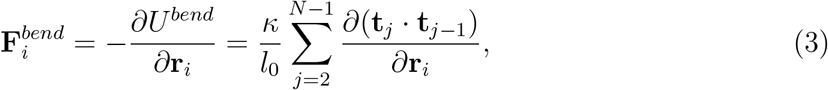

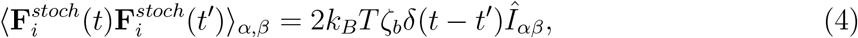

where

*N* is the number of beads per filament,

**t**_*j*_ = (**r**_*j*+1_ *−* **r**_*j*_)*/|***r**_*j*+1_ *−* **r**_*j*_| is the local unit tangent vector, *κ* = *k*_*B*_*Tl*_*p*_ is the flexural rigidity, *k*_*B*_ is Boltzmann’s constant, *T* is temperature, *l*_*p*_ is the persistence length of the filament, and 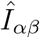 is the second order unit tensor. The spring constant between filament beads, *k*, is made smaller compared to the elongation modulus of actin filaments, in order to allow a longer simulation step [Laporte et al., 2012]. This filament model reproduces the correct tangent correlation function, relaxation dynamics and equi-partition of energy in thermal equilibrium.

The numerical values of the filament model parameters as well as other model parameters are shown in Table 1.

**Table 1:** Reference Parameter Values Used in Simulations

| Parameter | Description | value |
| --- | --- | --- |
| $R_{conf}$ | Confinement radius | $1.25 - 6 \mu m$ |
| $C_{actin}$ | Actin total concentration | $2 - 12 \mu M$ |
| $T$ | Temperature | 300 K |
| $l_p$ | Persistence length of actin filament | $17 \mu m$ |
| $\eta$ | Cytoplasmic viscosity (350 times larger than water) | $0.301 pN/\mu m^2 s$ |
| $k_+$ | Actin polymerization rate constant | $10 / \mu M / s$ |
| $l_0$ | Filament model segment equilibrium length | $0.1 \mu m$ |
| $k$ | Spring constant between filament beads | $100 pN/\mu m$ |
| $k_{atr}$ | Attraction force spring constant | $0.5 - 8 pN/\mu m$ |
| $r_{atr}$ | Attraction force range | $0.04 - 0.1 \mu m$ |
| $k_{srf}$ | Surface attraction force spring constant | $3 pN/\mu m$ |
| $r_{srf}$ | Surface attraction force range | $0.06 \mu m$ |
| $r_0$ | Equilibrium length for surface and attraction force | $0.012 \mu m$ |

A summary of the parameter values used in the different figures of the paper is shown in Table S1.

### 2.2 Attraction force between actin filaments

The attractive or crosslinking interaction between actin filaments is simulated by a shortrange, isotropic, attractive potential between filament beads (Fig. 1B-C). The force on bead bead *i* is within *r*_*atr*_ to bead *j* of the same or another filament is

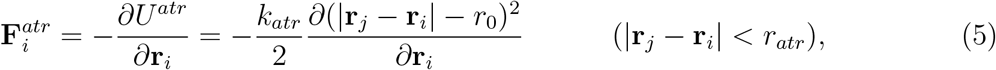

with spring constant *k*_*atr*_ and equilibrium length *r*_0_. Here *k*_*atr*_ and *r*_*atr*_ are effective parameters characterizing the depth and range of interaction between filament segments. This method can be used to study the non-specific attraction between actin filaments or the effective interaction through multiple crosslinker binding and unbinding between actin filament segments. A drawback of this method is that it does not explicitly account for the different molecular properties and concentration of crosslinkers.

### 2.3 Boundary conditions and interactions

We implemented three types of boundary conditions: I. Periodic boundary condition for bulk simulations; II. Repulsive hard wall, represented by a constant force of magnitude 1 *pN* normal to the cell boundary exerted to every bead crossing it; III. Short-range attraction near a spherical hard wall. For the latter case, we introduced an additional short-range surface spring force to each bead:

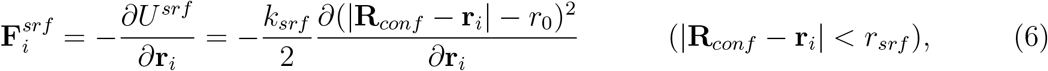

with spring constant *k*_*srf*_ and equilibrium length *r*_0_. Here *r*_*srf*_, is the distance from the surface in which beads can feel the surface force and **R**_*conf*_ is the location of the confining sphere boundary along the direction of **r**_*i*_.

### 2.4 Simulation of polymerization

Actin polymerization in vitro is typically initiated by addition of salts. This starts a process of nucleation and elongation as the unpolymerized monomer pool is depleted, followed by a much longer process of end-subunit exchange and length distribution relaxation that is not considered here [Oosawa and Asakura, 1975]. The average filament length at the time of monomer depletion depends on the concentration of nucleated filaments. This is a parameter that can be tuned experimentally: addition of severing proteins or preformed filaments seeds can speed up the polymerization process and lead to shorter filaments; profilin, which suppresses nucleation while typically not significantly affecting the elongation rate [Courtemanche and Pollard, 2013], can lead to longer filaments.

For simplicity, we assume a fixed number of filament nuclei at the start of the simulation. We vary the nuclei concentration to explore the dependence on final filament length as well as approximately match the experimental polymerization curves (see Results section). Polymerization of actin filaments is simulated as elongation of the first segment of the semiflexible polymer chain. As soon as the segment reaches twice the size of spring equilibrium length *l*_0_, a new bead is introduced to the polymer chain. The polymerization rate per filament, *k*_+_*C*(*t*) (in subunits per second), is proportional to the remaining bulk monomer concentration *C*(*t*) and the barbed end polymerization rate constant *k*_+_. For a fixed filament concentration, *C*(*t*) is a predefined function of time, decaying according to

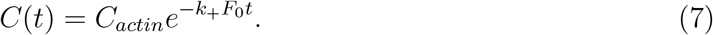

Here *F*_0_ is the filament nuclei concentration and *C*_*actin*_ is the initial bulk actin concentration. In the simulations all filaments reach the same length.

In the simulations we do not account for end-to-end annealing of actin filaments or filament polydispersity. We also neglect depolymerization from the barbed as well as polymerization and depolymerization from the slower pointed end.

Since typically *C*_*actin*_ is much larger than the critical concentration, the monomer concentration in Eq. 7 is assumed to decay to zero.

Even though the kinetics of Eq. 7 lack a nucleation phase, the timescale over which the monomers get depleted from the bulk is comparable to the experimental timescale measured in [Miyazaki et al., 2015] when the filament seed concentration is adjusted to provide a final filament length comparable to the average filament length measured in [Miyazaki et al., 2015] (Fig. S1).

### 2.5 Effect of excluded volume interactions

To check the effect of excluded volume repulsion between actin filaments at the low actin concentrations used in this work, simulations were performed following a method by Kim et al. [Kim et al., 2009]. Our implementation was for testing purposes and we did not use a more complex rigorous expression for excluded volume by Popov et al. [Popov et al., 2016]. The repulsive force between two filament segments is exerted along the direction of the line joining points *a, b* in each segment that are separated by the smallest distance, *r*_*ab*_. This defines a force *F*^*xv*^ = *k*_*xv*_(*r*_*ab*_ − *l*_*xv*_) when *r*_*ab*_ < *l*_*xv*_. Here *k*_*xv*_ = 1690 *pN/µm* and *l*_*xv*_ = 10.2 *nm* is of order the actin filament diameter. The repulsive force is distributed to the beads defining the segment end points in proportion to their distance to point *a* or *b* belonging to that segment [Kim et al., 2009]. A smaller time step, *dt* = 0.5× 10^*−*4^ *s* was required for numerical stability. We checked using the method of Sirk et al. [Sirk et al., 2012] that such a time step eliminated topological violations throughout the simulation.

A comparison of simulation results with and without excluded interaction is shown in Fig. S2. Addition of excluded volume repulsion results in somewhat thicker and less curved bundles. As an added repulsive interaction, excluded volume repulsion also leads to a small shift of the phase boundaries between bundled and unbundled phases described below. The overall effect of excluded volume repulsion is however relatively small. Thus, for numerical efficiency purposes, excluded volume interactions among filaments and cross-linker proteins is neglected in the rest of the paper.

### 2.6 Planar order parameter, radial density, and bundle thickness

We define a “planar order parameter” to quantify the degree with which the resulting network lies on a plane, as would be the case of a ring structure. This parameter measures the alignment of the filament segment binormal vectors, 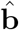 and it is the highest eigenvalue of tensor **Q** [Saupe, 1968]:

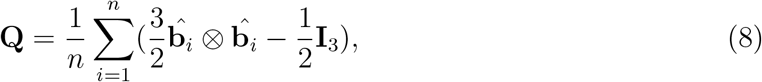

where *n* is number of filament segments (i.e. the filament segment between neighboring beads along a filament). This is a number between 0 and 1, with 0 representing random orientation and becoming 1 when all binormal vectors align (thus filaments are on a plane or parallel planes).

The normalized distribution of actin beads along the radial direction under spherical confinement is calculated by

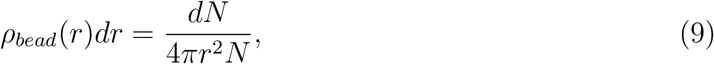

where *dN* is the number of filament beads within *dr* and *N* the total number of beads.

To calculate average properties, we repeated simulations with identical parameter values several times. The average radius of the configuration was calculated as follows:

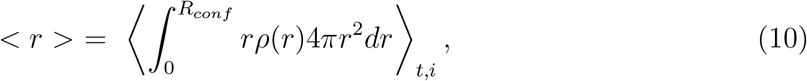

where *i* labels different simulation runs and the average over time *t* is specified below (typically performed over several time points during the second half of the simulation, 1000 - 1500 *s*). The number of runs (range of *i*), which varied depending on the accuracy required, is indicated in the figures. A similar averaging was performed for the average planar order parameter. The probability of ring formation was calculated as the fraction of times the system formed a closed circular loop that was stable through the end of the simulation.

The number of actin filaments in a bundle was determined by scanning along beads of each filament, finding the plane perpendicular to the filament at that point and calculating the number of beads of other filaments that are within 0.01 *µm* on either side of the plane and not further than 0.2 *µm* from the bead.

### 2.7 Simulation implementation

We developed Java code for polymerizing semiflexible filament simulations, extending and parallelizing methods of prior works [Tang et al., 2014, Bidone et al., 2014]. A typical run time over 1500 s for 5 *µM* actin in a sphere of radius 2.5 *µm* using three cores on XSEDE’s Comet (Intel Xeon E5-2680v3 2.5 GHz) was 48 hours.

## 3 Results

### 3.1 Actin distribution in confinement depends on both filament length and attraction strength

The schematic of our Brownian dynamics filament model and the short range attraction potential between filament beads is illustrated in Fig. 1A-B. The graph of Fig. 1C shows how increasing spring constant *k*_*atr*_ leads to a deeper attraction well. Since this effective potential may represent the effect of multiple crosslink binding and unbinding between filaments, a deeper well may be considered to represent the effect of higher crosslink concentration and/or less dynamic crosslinks that have longer lifetimes. We also explore the effect of the value of the interaction range *r*_*atr*_, up to the distance between two filament beads (*l*_0_ = 0.1 *µm*), even though *l*_0_ may be large compared to the typical size of actin filament crosslinkers or the range of depletion interactions. A larger value of *r*_*atr*_ can be considered to represent a larger crosslinker concentration (larger chance for two filament segments to interact) and/or larger crosslinker physical size. Large values of *r*_*atr*_ also allow easier filament sliding along actin bundles as described below.

An example of a simulation with 140 filaments polymerizing and associating with each other and forming a ring within a sphere of radius *R*_*conf*_ = 2.5 *µm* is shown in Fig. 1D. Unless otherwise indicated, here and below, a repulsive hard wall boundary is assumed. In our simulations, the filament elongation rate adjusts to match the calculated reduction in the available monomer pool. By the end of the simulation each filament reached a final length of 3.8 *µm*.

Actin filament association and crosslinking results in stiff bundles, the effective persistence length of which increases with bundle thickness, depending on depletant concentration and the type of crosslinker [Claessens et al., 2006a, Takatsuki et al., 2014]. Our simple coarse-grained representation of crosslinking reproduces qualitatively this stiffening effect, for example the persistence length of a bundle of five actin filaments increases to about 80 *µm* at sufficiently high crosslinking spring constant *k*_*atr*_ for *r*_*atr*_ = 0.06*µm* (Fig. S3). For *k_atr_ <* 1.5*pN/µm* the attraction is not strong enough to form a bundle and the persistence length is equal to the single filament persistence length (17*µm*). Increasing *k*_*atr*_ above 1.5 *pN/µm* leads to an increase in the bundle persistence length up to 80*µm*. This increase is comparable to studies of crosslinking by a flexible linker such as *α*-actinin and plastin in 1:50 crosslinker:actin concentration ratios [Claessens et al., 2006a, Heussinger et al., 2007].

To show how confinement influences filament shape and distribution, we first performed simulations of spherically confined non-interacting filaments (*k*_*atr*_ = 0) of fixed length. Filaments of length *l*_*fil*_ much shorter than the confinement radius *R*_*conf*_ are distributed with an almost uniform *ρ_bead_*(*r*) distribution within the center of the sphere, with a reduced concentration near the boundary where orientational restrictions result in entropy loss (Fig. 2A,B). Since the persistence length of these short filaments is much longer than their contour length, their shapes are approximately straight lines. Filaments of length comparable to the diameter, *l*_*fil*_ = 1.8*R*_*conf*_ remain close to the center where they can also remain straight along the diameter of the sphere (Fig. 2A). As a result, *ρ_bead_*(*r*) peaks in the center of the sphere (Fig. 2B). Increasing filament length to *l*_*fil*_ = 8*R*_*conf*_ causes the filaments to bend and adopt the shape of the confining sphere [Morrison and Thirumalai, 2009], with *ρ_bead_*(*r*) peaking close to the confining boundary surface. These results are in qualitative agreement with [Gârlea, 2015] who used a different numerical method.

**Figure 2:**
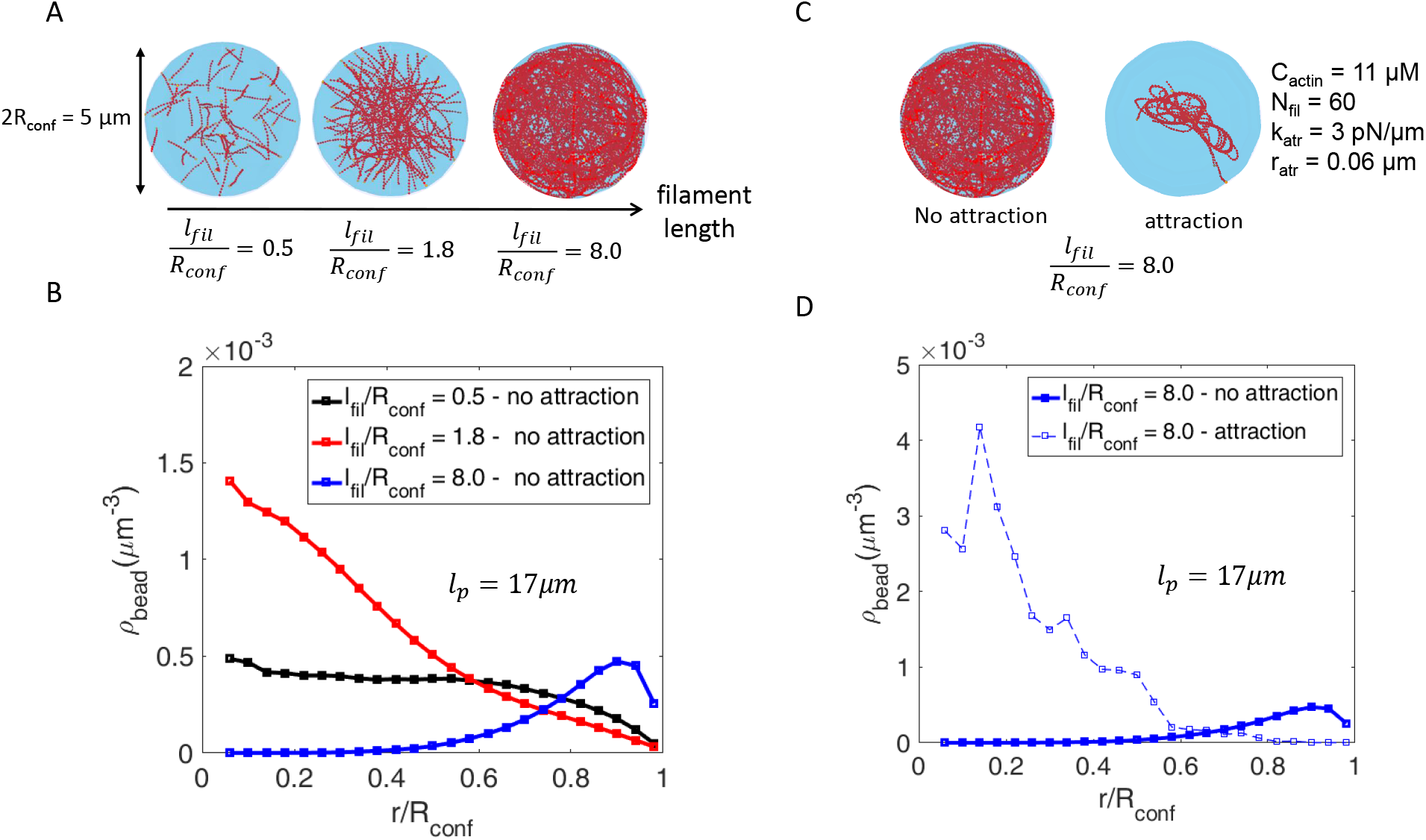
Attraction influences configuration of confined actin filaments. (A) Snapshots of simulations with 60 non-interacting filaments after 1500 *s* in a sphere of radius *R*_*conf*_ = 2.5 *µm*. The three cases show filaments of length *l*_*fil*_ = 1.2, 4.5, and 20 *µm*. (B) Radial bead density plotted as function of distance from sphere center for the three cases in panel A. (C) Snapshots of simulations at 1500 *s*, for *R*_*conf*_ = 2.5 *µm* and 60 filaments with *l*_*fil*_ = 20 *µm* without and with attraction (*k*_*atr*_ = 3 *pN/µm, r_atr_* = 0.06 *µm*). Here the initial concentration of actin monomers was 11 *µM*. (D) Radial bead density plotted as function of distance from sphere center for the two cases of panel C. In panels B and D the average is calculated by sampling beads in all 60 filaments from 500 *s* to 1500 *s* every 5 *s*. The curves in panels B and D become noisier close to *r* = 0 (and thus not plotted very close to 0) due to fluctuations in the number of beads over a small volume, as well as the irregular structure of the collapsed filaments in panel C.

Attractive interactions between filaments are also important in filament distribution. As a direct example, we consider the same case of long filaments *l*_*fil*_ = 8*R*_*conf*_ as in Fig. 2A and turn on an attractive force. Instead of bundling the filaments into a ring along the sphere, as seen in Fig. 2C,D, this attraction caused the filaments to collapse into the center of the sphere where they can gain energy by interacting with each other. This suggests that the ring formation observed in Fig. 1D requires tuning of filament length, confining radius and attractive strength. We explore this dependence systematically below after first comparing to the very different network structures that occur in a bulk environment.

### 3.2 Actin network formation in the bulk

To compare and contrast confined versus bulk cases, we simulated actin polymerization in the bulk using a 5 *µm* box with periodic boundary conditions (Fig. 3). Here and in most simulations below we use *C*_*actin*_ = 5.0 *µM*, a typical bulk actin concentration used in experiments in vitro. This concentration is also chosen to be between the two cases of 2 and 10 *µM* studied in [Miyazaki et al., 2015]. For the bulk simulations of Fig. 3 we kept the attraction range to *r*_*atr*_ = 0.06 *µm* and varied the strength of crosslinking by changing the crosslinking spring constant *k*_*atr*_ = 0.3 *−* 3 *pN/µm*. We adjusted the initial filament nuclei concentration to get a final filament length *l*_*fil*_ = 3.8 *µm* at long times.

**Figure 3:**
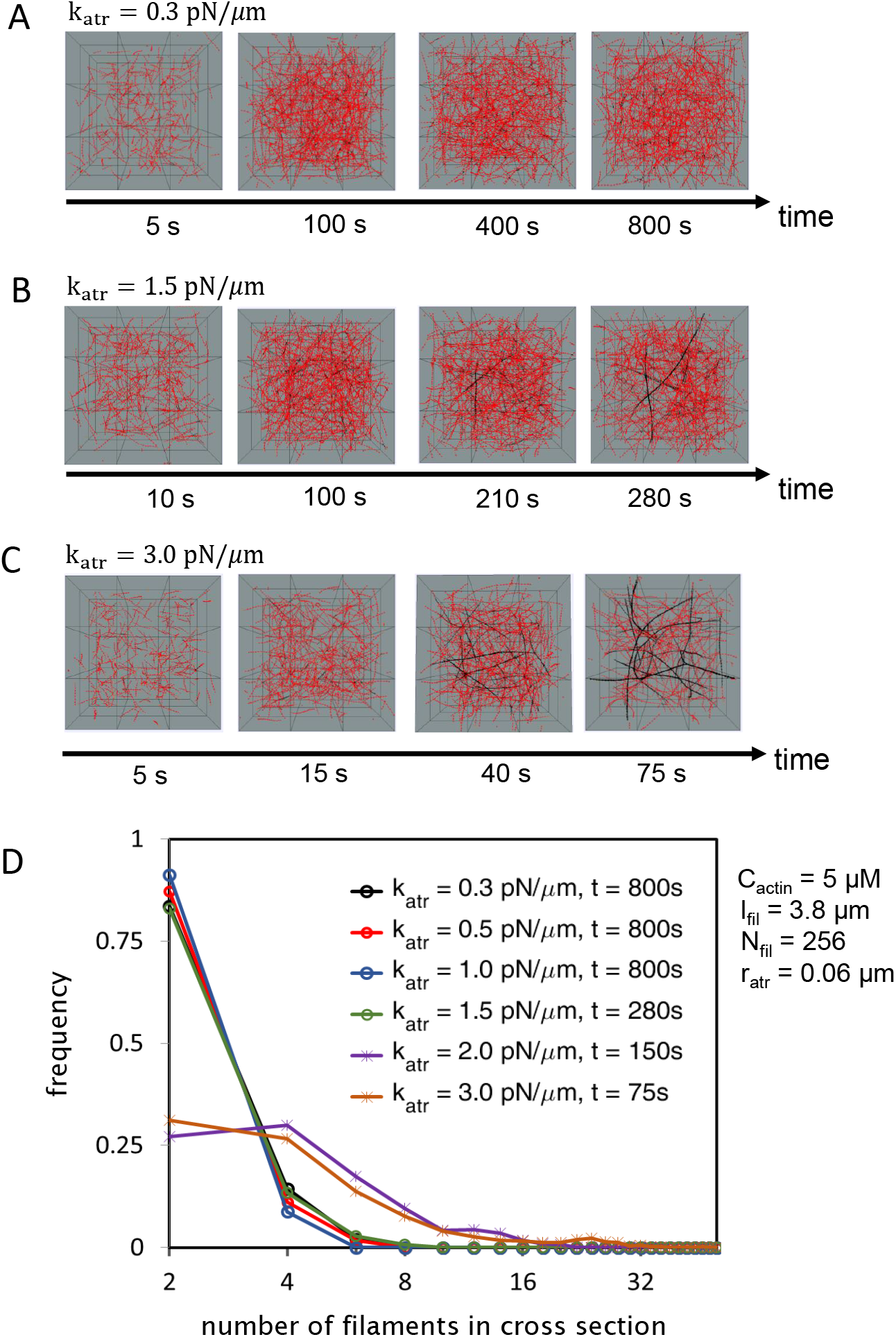
Simulations of polymerizing and associating actin filaments in the bulk. (A-C) Snapshots of simulation for a square box of size 5*µm*, periodic boundary conditions, and 265 filaments polymerizing from an initial 5 *µM* monomer concentration. Filaments reach length *l*_*fil*_ = 3.8 *µm*. Beads in range (not in range) to interact with neighboring beads are shown in black (red). In all cases *r*_*atr*_ = 0.06 *µm*. As *k*_*atr*_ is varied, a transition from an unbundled phase to a to network of bundles occurs at around *k*_*atr*_ = 1.5 *pN/µm*. Simulations in panels B and C are terminated before the network starts to deform in an unphysical manner as a result of the periodic boundary conditions. (D) Normalized distribution of thickness (measured in number of filaments, see Model and Methods section) of actin filament bundles for various values of *k*_*atr*_, at the indicated times, which correspond to the times at which the simulations were terminated. The fraction of beads interacting with other beads is 0.035, 0.027, 0.028, 0.035, 0.526, 0.527 for *k*_*atr*_ = 0.3, 0.5, 1.0, 1.5, 2.0, and 3.0 *pN/µm* respectively.

We observed unbundled and bundled phases. For weak crosslinking, *k*_*atr*_ ≤ 1.0 *pN/µm*, the attraction is not strong enough to stabilize bundles (illustrated by black color in Fig. 3) against thermal fluctuations. Thus the system evolves into an unbundled phase with the thickness of any transient bundles (mostly dilute contact points) being typically 2 filaments (Fig. 3A,D). As the attraction strength is increased to *k*_*atr*_ = 1.5 *pN/µm*, thin long bundles form (Fig. 3B,D). For even higher strength, *k_atr_ >* 2 *pN/µm*, a network of thick bundles develops quickly (Fig. 3C,D). As will also be discussed below, very little filament sliding within bundles and very little unbundling occurs for these high *k*_*atr*_ values and *r*_*atr*_ = 0.06 *µm*. The simulations in Fig. 3B,C were terminated once the bundle network spanned the simulation box size: periodic boundary conditions allow the network to slide along the boundary and collapse into a less dense structure in ways that would not be possible in a real 3D system. Despite this technical limitation, these simulations show how actin filaments at *µM* concentrations in a bulk undergo an abrupt transition to a bundled network, as predicted theoretically [Zilman and Safran, 2003], without forming loop or ring structures.

### 3.3 Associating actin filaments in confinement: effect of filament length and confinement radius

As discussed in the preceding sections, both confinement and attractive potential can influence the behaviour of the system. Furthermore, the behaviour of filaments in confinement depends on the final filament length relative to the size of the confining space (Fig. 2B). This observation leads us to study the effect of three factors: final filament length *l*_*fil*_, attraction force, and confining diameter 2*R*_*conf*_. First, we investigate three different filament length regimes: *l_fil_ <* 2*R*_*conf*_, *l_fil_ ≈* 2*R*_*conf*_ and *l_fil_ ≫* 2*R*_*conf*_, with constant confinement radius *R*_*conf*_ = 2.5*µm* and concentration *C*_*actin*_ = 5 *µM*. We explore a range of attraction strength by varying both parameters describing this interaction, *k*_*atr*_ and *r*_*atr*_, in Figs. 4, 5 and 6. The effect of reducing the confinement radius to *R*_*conf*_ = 2*µm* is shown in Fig. S4-S6. The final length of the filament in these simulations was changed by changing the initial filament nuclei concentration. To quantify the resulting actin organization we used the average radius, the planar order parameter describing the organization along a plane, and the probability of ring formation (see Model section). In all cases we evolved the system up to 1500 *s* and quantified the long-time behavior in the 1000-1500 *s* interval. In this section we kept the confining boundary condition to be a repulsive hard wall. The results are summarized in Fig. 7.

**Figure 4:**
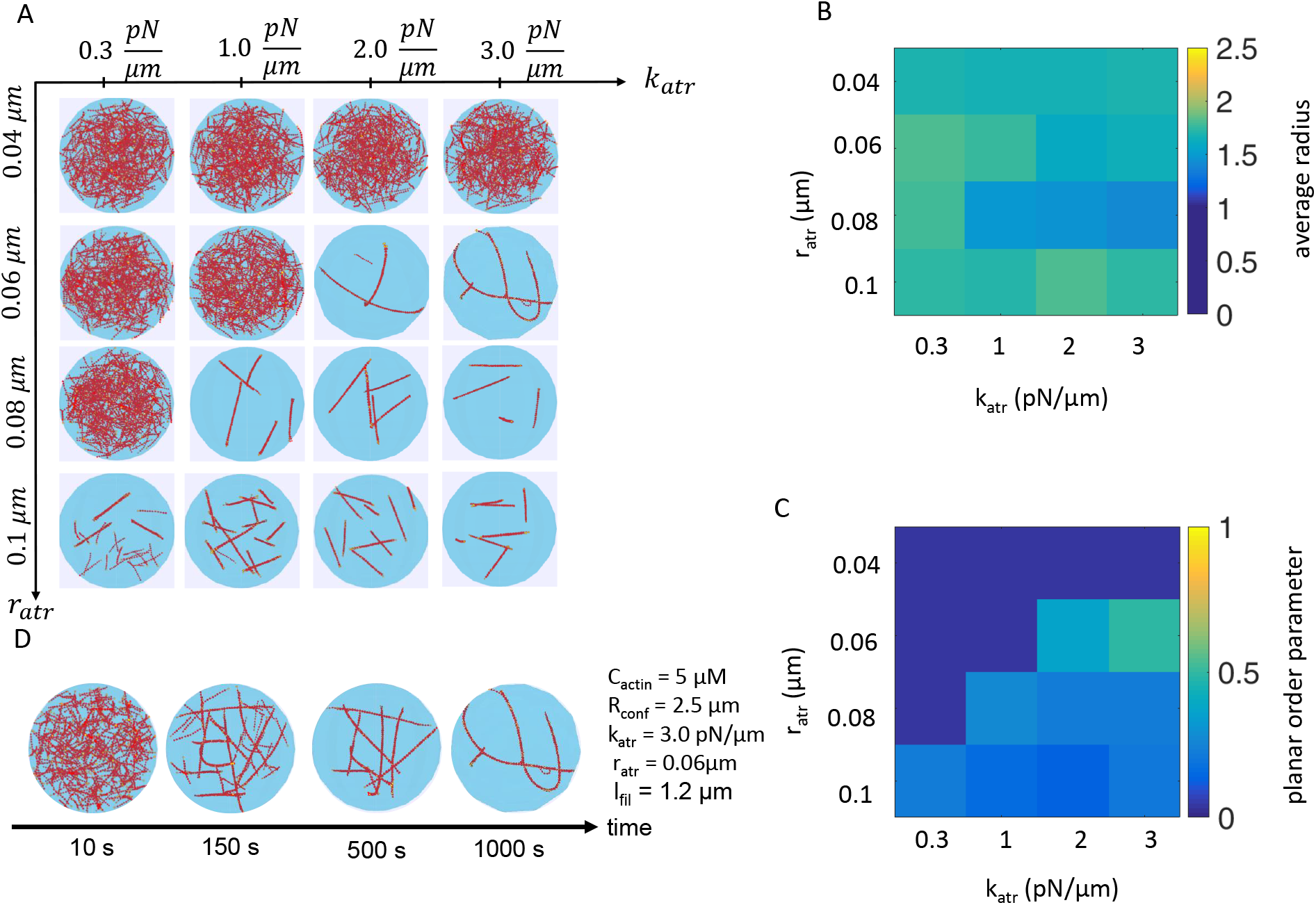
Actin network morphology for filaments that elongate to lengths shorter than the diameter of the confinement. The final configuration depends on the strength of attraction *k*_*atr*_ and range of interaction *r*_*atr*_. (A) Snapshots of simulation at 1500 *s* for *C*_*actin*_ = 5 *µM*, *R*_*conf*_ = 2.5 *µm*, 420 filaments, and final filament length *l*_*fil*_ = 1.2 *µm* for four values of *r*_*atr*_ and four values of *k*_*atr*_ as indicated. (B) Average radius, sampled from 1000 to 1500 *s* every 10 *s*, as a function of *r*_*atr*_ and *k*_*atr*_, averaged over 3 separate runs. (C) Average planar order parameter, sampled as in panel B. (D) Snapshots of the case of *k*_*atr*_ = 3.0 *pN/µm* and *r*_*atr*_ = 0.06 *µm* over time.

**Figure 5:**
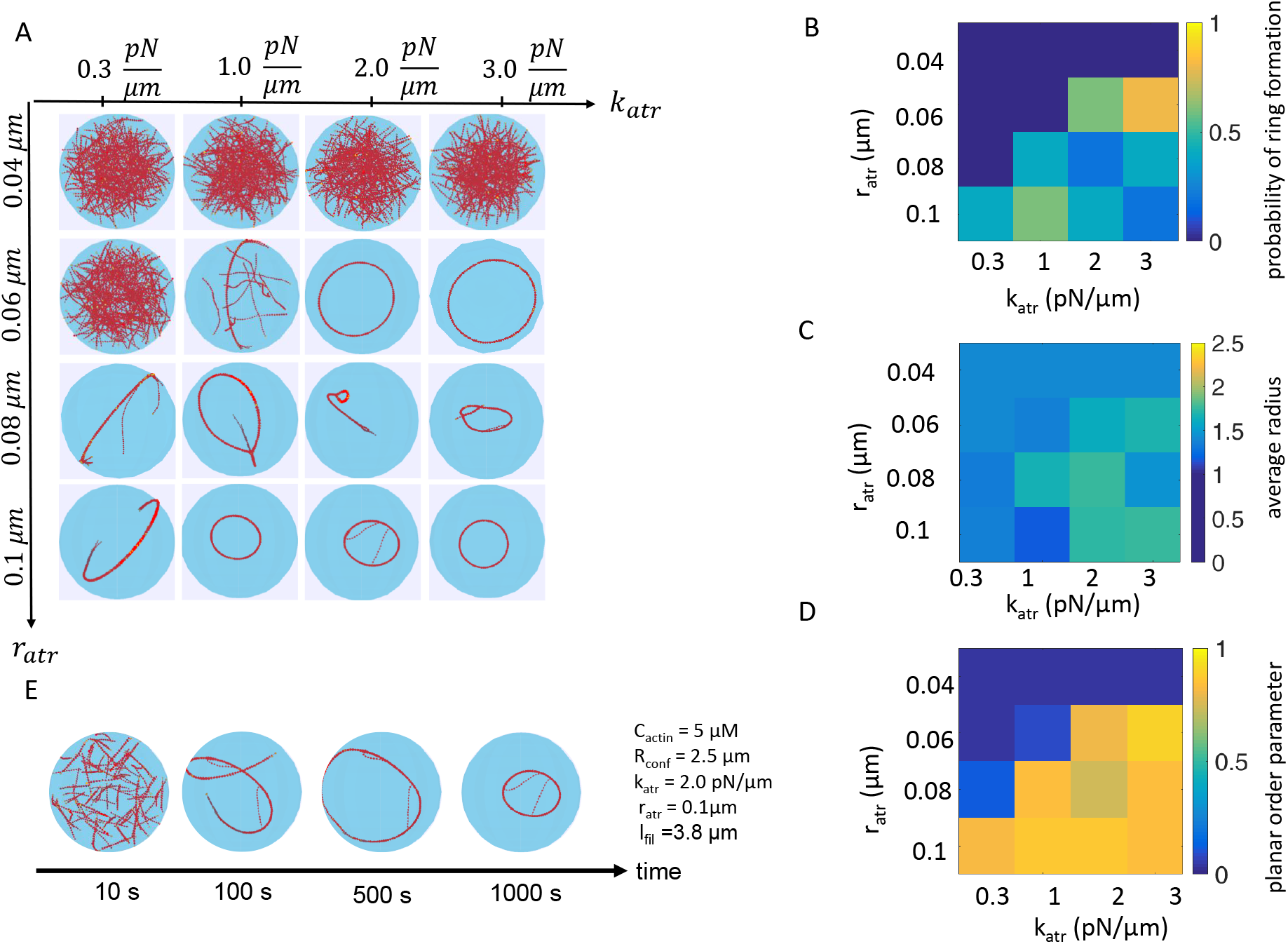
Actin network morphology for filaments that elongate to lengths comparable to the diameter of the confinement. Similar to Fig. 4, with the addition of panel B showing the probability of ring formation. (A) Snapshots of simulation at 1500 *s* for *C*_*actin*_ = 5 *µM* and *R*_*conf*_ = 2.5 *µm*, and 140 filaments reaching *l*_*fil*_ = 3.8 *µm*. Snapshots show representative examples; results can vary in each run, with rings, open bundles, or both occuring with different probability for the same parameter values. (B) Plot of probability of ring formation for 5 runs. (C) Average radius, sampled from 1000 to 1500 *s* every 10 *s* for 5 runs. (C) As panel C, for planar order parameter. (E) Snapshots of the case *k*_*atr*_ = 2 *pN/µm* and *r*_*atr*_ = 0.1 *µm* showing ring formation and shrinkage.

**Figure 6:**
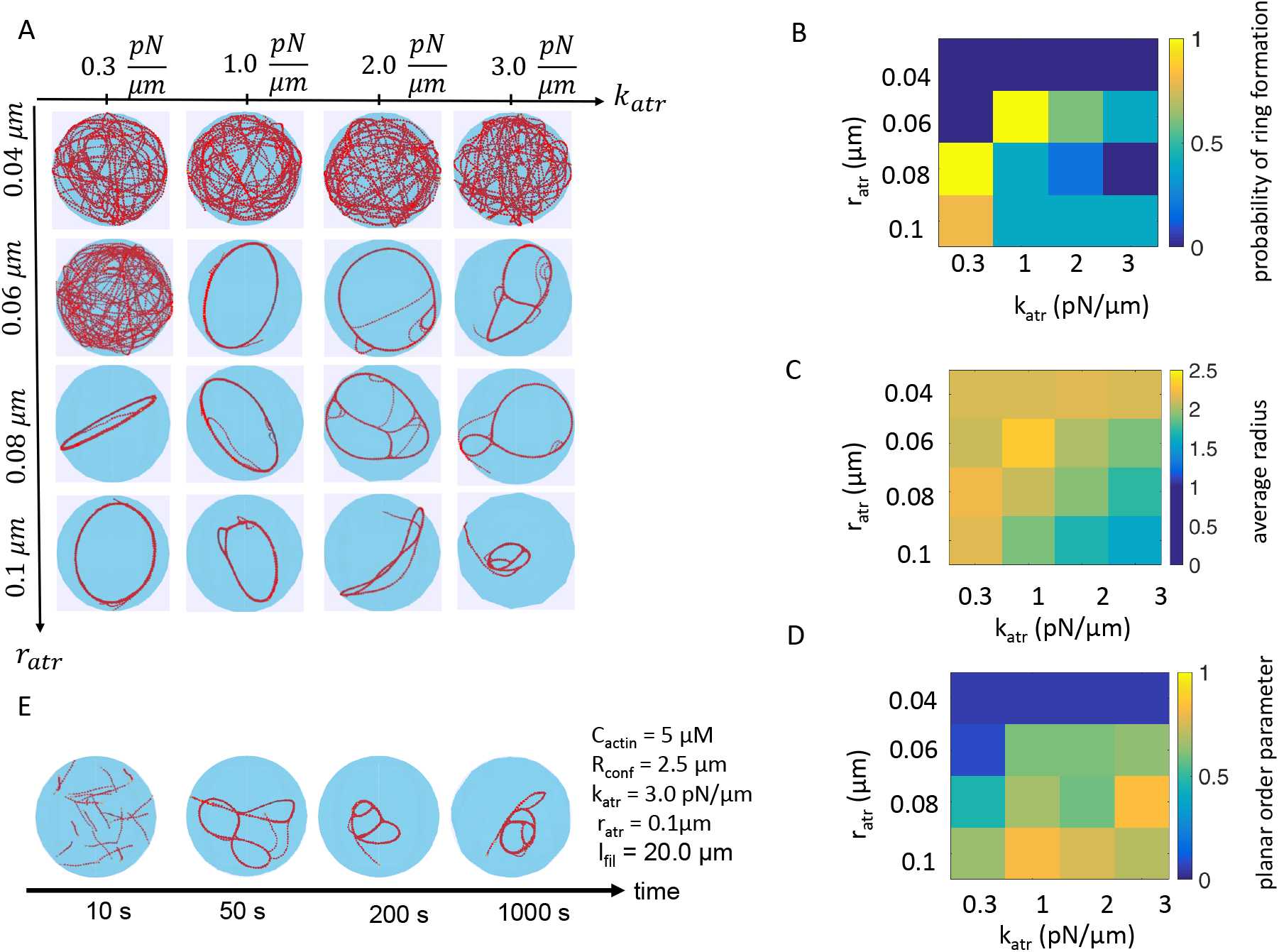
Actin network morphology for filaments that elongate to lengths longer than the diameter of the confinement. (A)-(D) Results for *C*_*actin*_ = 5 *µM*, *R*_*conf*_ = 2.5 *µm* presented in the same way as in Fig. 5 but for 26 filaments reaching *l*_*fil*_ = 20 *µm*. Snapshots show representative examples; results can vary in each run. The averages in panels B-D are over 5 runs. (E) Snapshots of the case of *k*_*atr*_ = 3 *pN/µm* and *r*_*atr*_ = 0.1 *µm* showing the development of an aggregated structure over time.

**Figure 7:**
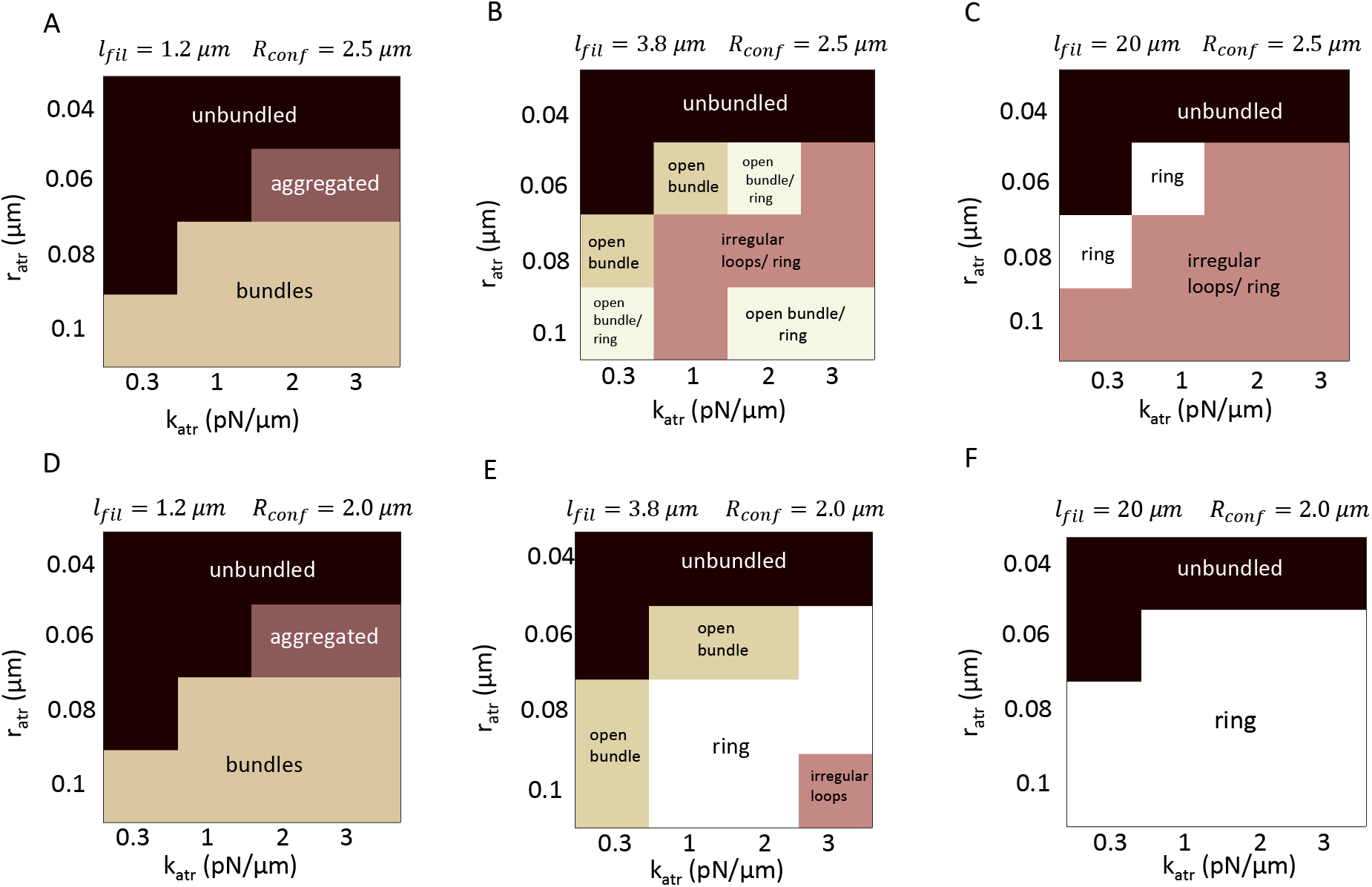
Summary diagrams of confined actin structures, as a function of the strength of attraction *k*_*atr*_, range of interaction *r*_*atr*_, for varying final filament length and confining radius. All cases are for *C*_*actin*_ = 5*µM*. (A)-(C) Results for *R*_*conf*_ = 2.5 *µm* from Figs. 4-6. (D)-(F) Results for *R*_*conf*_ = 2.0 *µm* from Figs. S4-S6. Labels indicate the dominant structures for each parameter value.

#### 3.3.1 Short filaments, *l_fil_ <* 2*R*_*conf*_

For *l*_*fil*_ = 1.2 *µm*, polymerization is nearly complete at *t*_*poly*_ = 60 *s*. The system explores three main configurations: unbundled, disconnected short bundles, and aggregated, while ring formation was not observed (Fig. 4A and Fig. 7A). The unbundled configuration occurs for weak attractive force (small *r*_*atr*_ or *k*_*atr*_), where the filaments show a nearly uniform distribution and the planar order parameter is close to zero (Fig. 4B and C). Disconnected short bundles occur for *r*_*atr*_ ≥ 0.08 *µm* because this large value of *r*_*atr*_ allows sliding of filaments relative to each other within a bundle: the range of attraction between beads is larger than the distance between two beads along the filament, leading to a relatively smooth potential as a function of sliding distance along the bundle axis. Thus, interacting filaments compact to maximize binding energy. The resulting disconnected bundles slowly coalesce with each other in a process that is not complete by 1500 *s*. These bundles do not develop into tactoidal droplets as this would require higher actin concentration (*>*100 *µM*) [Oakes et al., 2007]. (We note that a tactoidal shape starts to appear in simulations with excluded volume interactions, Fig. S2).

An interesting case occurs for *r*_*atr*_ = 0.06 *µm* and high *k*_*atr*_, where short filaments come together to make an “aggregated” structure containing bundles of length longer than that of individual filaments (Movie S1). This phenomenon is similar to the network formation process in the absence of confinement in Fig. 3D that used the same *r*_*atr*_ value. In this case, filament sliding within a bundle occurs very slowly, as a result of the strong and disconnected crosslinks among filaments. Thus, the resulting aggregate is in a kinetically trapped state that has a degree of planarity (Fig. 4C) but does not come together into a ring because bundles start to form at multiple random orientations, as seen in the snapshots of Fig. 4D and Movie S1. We would expect such structures to form in the presence of effectively irreversible crosslinking, which may occur as a result of long-lived crosslink lifetimes or very strong depletion forces [Ward et al., 2015, Rückerl et al., 2017].

The qualitative phase diagram of Fig. 7A does not change in simulations with a smaller confining radius but same filament length and actin concentration (*R*_*conf*_ = 2.0*µm*, Fig. S4 and Fig. 7D).

In summary, rings do not form in this case of filaments short compared to the confining radius. By increasing parameters *r*_*atr*_ or *k*_*atr*_ we can induce the bundling transition. For *r*_*atr*_ = 0.06 *µm* and *k_atr_ >* 2 *pN/µm* the simulations show the effect of kinetic trapping in bundles as might occur for crosslinking by long-lived crosslinkers. For *r_atr_ >* 0.08 *µm* we have filament sliding within bundles as might occur for intermediate depletion interaction strength [Ward et al., 2015] (even though the value of the range of depletion interaction would be smaller in magnitude compared to our model in an actual physical system) or sufficiently dynamic crosslinking.

#### 3.3.2 Intermediate length filaments, *l_fil_ ≈* 2*R*_*conf*_

To study the case of *l_fil_ ≈* 2*R*_*conf*_, we tuned the number of filament nuclei such that the final filament length is *l*_*fil*_ = 3.8*µm*. In this case polymerization is nearly complete after *t*_*poly*_ = 200 *s*. We found four types of different architectures (Fig. 5A and Fig. 7B). As for the case of short filaments, an unbundled phase is found for weak attractive interactions (small enough *r*_*atr*_ and *k*_*atr*_). The size of the unbundled region shrank compared to Fig. 4A as a result of the increased filament length. The three other bundled configurations include “open bundle”, ring, and irregular looped structures (Fig. 5A, Fig. 7B, Movies S2 and S3). By open bundle we mean a curved bundle that does not close up into a ring. The boundaries between those regions were not sharp: different outcomes (such as open bundle, ring, or both together) could result for the same parameter values on different runs.

The maximum probability of ring formation within the range of attraction parameters explored in this figure was 80% at *k*_*atr*_ = 3.0 *pN/µm*, *r*_*atr*_ = 0.06 *µm* (Fig. 5B, out of five runs). Interestingly, even though these two parameter values lead to kinetically trapped states for shorter filament lengths (Fig. 4A), here, a ring forms robustly, as a result of the smaller number of longer filaments (less possible alignment orientations) and confinement-induced bending of attracted filaments that elongate without significant sliding along the developing and looping bundle. The resulting ring goes close to the boundary (Fig. 5C) and has high value of the planar order parameter (Fig. 5D).

The configurations of filaments in “open bundle”, ring, and irregular looped structures (that resemble the lollipop shapes of individual self-associating filaments [Lau et al., 2009] or unequal double rings) occur in the region of phase space where filament sliding along bundles can occur more easily (*r*_*atr*_ ≥ 0.08*µm*). This reflects the existence of multiple topologically-different free energy minima in the system (as can also occur in the collapse of single self-attractive filaments [Ou and Muthukumar, 2005, Lappala and Terentjev, 2013]). Confinement enables the system to explore and evolve towards these minima as follows. When the kinetics of bundle assembly lead to a strand that is long enough to make a closed loop as in Fig. 5E, the system proceeds to lower its free energy through ring constriction that maximizes contacts at the expense of bending energy. The radius of these rings can become smaller than the confinement radius (average radius smaller than *R*_*conf*_, see Fig. 5C). However, if the developing bundle does not elongate long enough to be able to fold into a ring (because of the kinetics of association in the presence of compaction by filament sliding within bundles) or in case the ring opens up, the free energy is lowered through further bundle compaction and further opening up of the gap.

Interestingly, reducing the confining radius to *R*_*conf*_ = 2.0*µm* greatly promotes ring formation when we keep the concentration and final filament length fixed (Fig. 7E and Fig. S5). There are two main factors responsible for this change. First, the smaller radius imposes greater bending, facilitating loop closure. Second, decreasing the number of filaments from 140 to 69 leads to less variability in loop number and shape.

#### 3.3.3 Long filaments, *l_fil_ ≫* 2*R*_*conf*_

As a third case, we initialized an even smaller number of filaments such that the final length is *l*_*fil*_ = 20 *µm*. In this case polymerization is nearly complete by *t*_*poly*_ = 800 *s*. Since filaments are now long enough to loop back into themselves within the confining sphere, we don’t find any open bundles. The system shows three phases: unbundled, for weak interaction strength, and ring or irregular loops for larger interaction strength (Fig. 6A, Fig. 7C and Movie S4).

The irregular structures are once again indicative of distinct topological configurations enabled by confinement. The longer filament length in this case slows down loop shrinkage through filament sliding along bundles. However we do observe the average radius decreasing with increasing *k*_*atr*_ and *r*_*atr*_ (Fig. 6C), an effect that happens before the polymerization process is complete (Fig. 6E).

Ring formation is robust close to the boundary with the unbundled phase (Fig. 6B and Fig. 7C). These rings, which have nearly the largest possible radius, form near the end of the 1500 *s* simulation. This is the reason why the planar order parameter of Fig. 6D, which is measured in the whole interval 1000-1500 *s*, is less than unity. These rings should correspond to a thermal equilibrium configuration at a sweet spot where crosslinking is both strong enough to create a bundle but not as strong to force filament bending (for contact increase) and to avoid kinetic traps such as multi-loop formation. Decreasing the confining radius greatly promotes ring formation (Fig. S6 and Fig. 7F), an effect similar to the case of intermediate length filaments.

### 3.4 Comparison to prior experiments: effect of change in confinement radius, concentration, and confinement shape

Having studied how ring formation depends on several parameters in Section 3.3, we now proceed to test if we can interpret the experimental results of Miyazaki et al. [Miyazaki et al., 2015] who systematically varied droplet size, concentration and confinement shape. They studied actin ring formation and contraction in the presence of motor proteins (HMM and myosin V), *α*-actinin crosslinkers, methylcellulose depletion agent, and combination of them. Each one of these factors can act to promote or prevent ring formation in spherical confinement. Here we first focus on the experimental results in the presence of methylcellulose, the concentration of which was varied to optimize ring formation. The observed effects of motor proteins on ring assembly and constriction that are beyond the scope of our model are briefly discussed in the Discussion section.

#### 3.4.1 Confinement radius

Miyazaki et al. [Miyazaki et al., 2015] found that an increase in confinement droplet radius led to a decrease in the probability of ring formation for both 2 *µM* and 10 *µM* actin concentration. Indeed, in the previous section we also found that *R*_*conf*_ has an important impact on the likelihood of ring formation (Fig. 7). For a more detailed comparison, we kept the actin concentration to 5 *µM*, a value in between the two concentrations in Miyazaki et al.

We adjusted the fixed filament concentration to *F*_0_ = 2.2 *nM*, which gives a final filament length *l*_*fil*_ = 6.0 *µm*. This length is close to the measured average length of 4.6 *µm* of actin filaments in droplets at 10 *µM* concentration [Miyazaki et al., 2015]. This *F*_0_ value also captures the approximate polymerization kinetics with a polymerization curve in between the 2 and 10 *µM* experimental curves (Fig. S1) (even though our simulations do not include nucleation, filament severing, or polydispersity).

As our choice for association parameters, we examined two parameter sets (PS): PS1, *k*_*atr*_ = 0.3 *pN/µm, r_atr_* = 0.1 *µm* and PS2, *k*_*atr*_ = 3.0 *pN/µm, r_atr_* = 0.08 *µm*. These two sets of values were selected using the phase diagram of Fig. 5 and Fig. 7B, corresponding to the case of filament length comparable to confining diameter, similar to what happens in the smaller droplets in [Miyazaki et al., 2015]. More specifically, both parameter sets gave a probability of ring formation similar to the expected probability (as the experiment trend shows) for confining radius 2.5*µm* (Fig. 8C).

**Figure 8:**
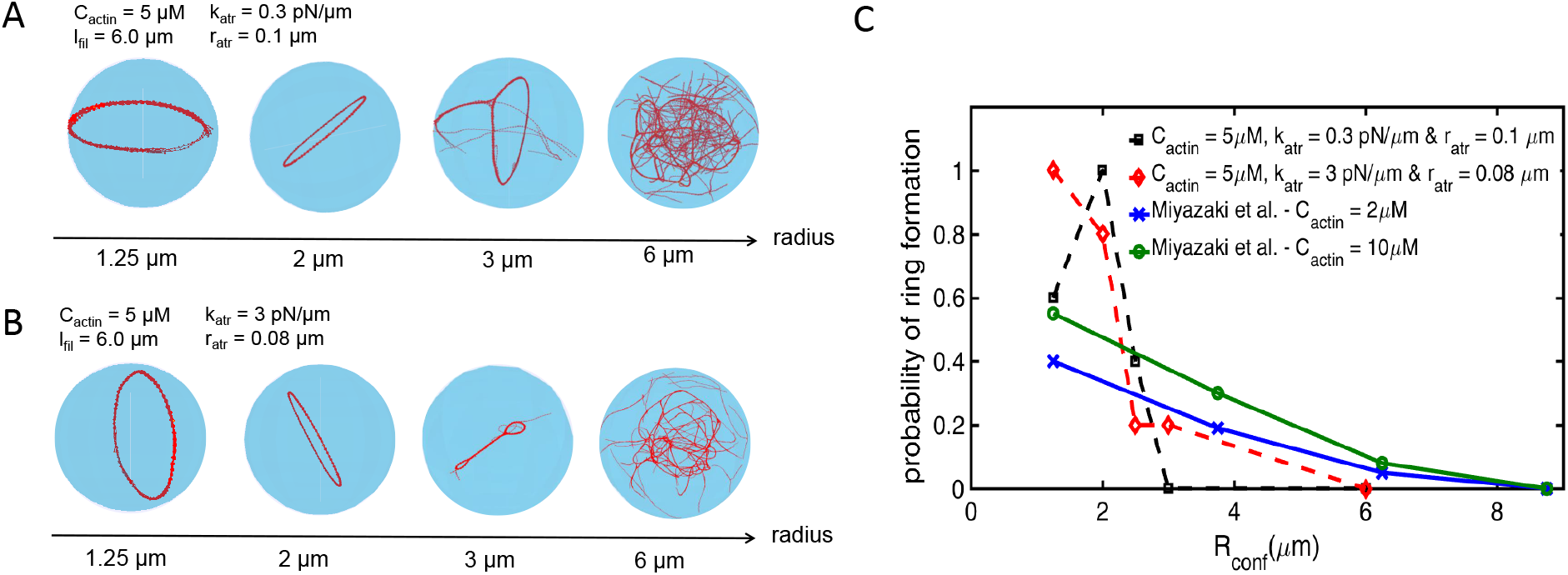
Probability of ring formation vs. size of the confinement. (A) Snapshots of simulation at 200 *s* for *C*_*actin*_ = 5*µM*, using parameter set PS1 (*k*_*atr*_ = 0.3 *pN/µm, r_atr_* = 0.01 *µm*) and varying confining radius *R*_*conf*_ as shown. The corresponding number of filaments in the four cases were 11, 45, 153, and 1227 such that the final filament length is always *l*_*fil*_ = 6 *µm*. (B) Same as panel A, for PS2 (*k*_*atr*_ = 3 *pN/µm, r_atr_* = 0.08 *µm*). (C) Probability of ring formation as function of confinement radius (out of 5 runs) for PS1 (dashed black line) and PS2 (dashed red line) compared to experiment data reproduced from Miyazaki et al. [Miyazaki et al., 2015] for *C*_*actin*_ = 2 *µM* (blue solid line) and *C*_*actin*_ = 10 *µM* (green solid line).

For both PS1 and PS2, the probability of ring formation shows an overall decreasing trend with increasing size of confinement, in rough qualitative agreement with the experimental observations (Fig. 8). The agreement is better for PS2 while PS1 shows a non-monotonic trend. This trend of PS1 where decreasing the radius down to *R*_*conf*_ = 1.25*µm* decreases ring formation probability was also seen in Fig. 7B and E and is due to the compaction through sliding of a smaller number of filaments into an open bundle. For large radii, as we have seen before, rings fail to form because of irregular loop structures. The latter start to resemble a bulk network at the largest radius in Fig. 8A,B.

Miyazaki et al. [Miyazaki et al., 2015] found that adding *α*-actinin to the methylcellulose and actin solution follows the same trend of decreasing ring formation probability versus increase in confinement radius. However the chance of ring formation decreased as *α*-actinin concentration increased. This demonstrates the fact that the type of attraction is important for ring formation.

Finally, we note that the above results are also generally consistent with Limozin and Sackmann [Limozin and Sackmann, 2002] who found that rings of actin (in 2-10 *µM* concentration) and *α*-actinin were formed in small vesicles with *R_conf_ <* 6*µm*. Large vesicles contained a spiderweb-like network similar to the largest spheres in the simulations of Fig 8. Decreasing the actin filament length by the addition of severing protein severin resulted in mobile assemblies of short bundles while some small vesicles (*R_conf_ <* 4.5 *µm*) contained rings [Limozin and Sackmann, 2002]. This observation is in agreement with simulations of bundle formation by short filaments in Fig. 3 and the requirement of long enough filaments for ring formation in simulations.

#### 3.4.2 Concentration

Next we explored the effect of concentration in order to interpret the concentration dependence of ring formation probability in the experiments (Fig. 8). We examined the same association parameters PS1 and PS2 and adjusted *F*_0_ for each concentration, assuming the final filament length is approximately the same in each case. Experimental measurement shows that bulk actin polymerization for 10 *µM* of actin was nearly complete in the first 10 minutes and for 2 *µM* in 60 minutes. Picking *F*_0_ to maintain final filament length approximately reproduces the time course of polymerization, except for the smallest concentrations (for *C*_*actin*_ = 2 *µM* we have *F*_0_ = 0.3 *nM* though a larger value would give a better fit to polymerization, Fig. S1).

We find that indeed, for both PS2 and for PS1 above 3 *µM*, the ring formation probability decreases with increasing concentration. High concentrations result in the formation of irregular structures such as multiple loops and lollipop shapes (Fig. 9). Rings do not form in case PS1 for the lower concentration (2 *µM*). In this case filament sliding leads to the formation of an open bundle, similar to the results for PS1 at low confining radius in Fig. 8A,C.

**Figure 9:**
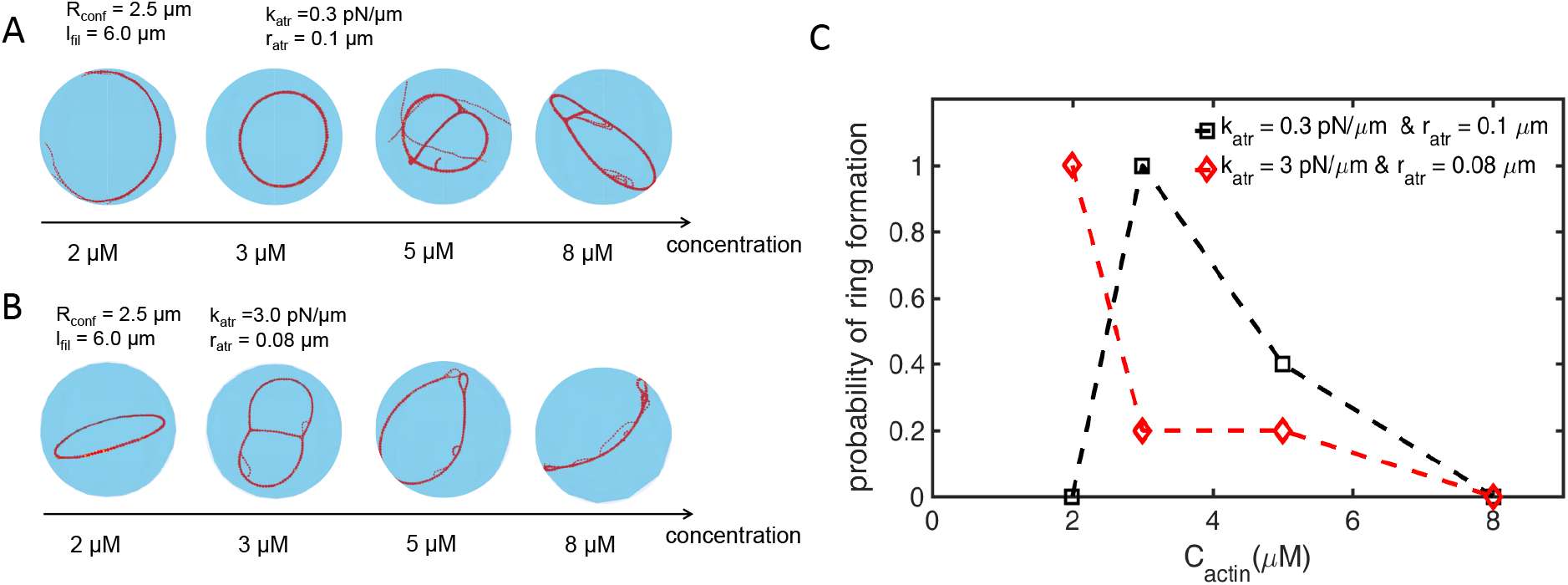
Probability of ring formation vs. actin concentration (A) Snapshots of simulation at 500 *s* for *R*_*conf*_ = 2.5*µm*, using parameter set PS1 (*k*_*atr*_ = 0.3 *pN/µm, r_atr_* = 0.1 *µm*) and varying concentration *C*_*actin*_ as shown. The corresponding number of filaments in the four cases were 36, 54, 89, and 142 such that the final filament length is always *l*_*fil*_ = 6 *µm*. (B) Same as panel A, for PS2 (*k*_*atr*_ = 3 *pN/µm, r_atr_* = 0.08 *µm*). (C) Probability of ring formation as function of actin concentration (out of 5 runs) for PS1 (black) and PS2 (red).

#### 3.4.3 Confinement geometry

We next simulated ring formation within oblate ellipsoids, to check if our results agree with the observation that the ring assembles at the plane perpendicular along the direction of droplet compression, where the curvature of the boundary is minimal [Miyazaki et al., 2015]. The results are shown in Fig. 10A,B for *C*_*actin*_ = 5*µM* and final filament length *l*_*fil*_ = 6*µm*, using PS1 for attraction. In these simulations the sphere had radius 2 *µm* and the oblate ellipsoid’s semi-minor axis was varied, keeping the volume fixed (33.5 *µm*^3^). Rings formed in all cases.

**Figure 10:**
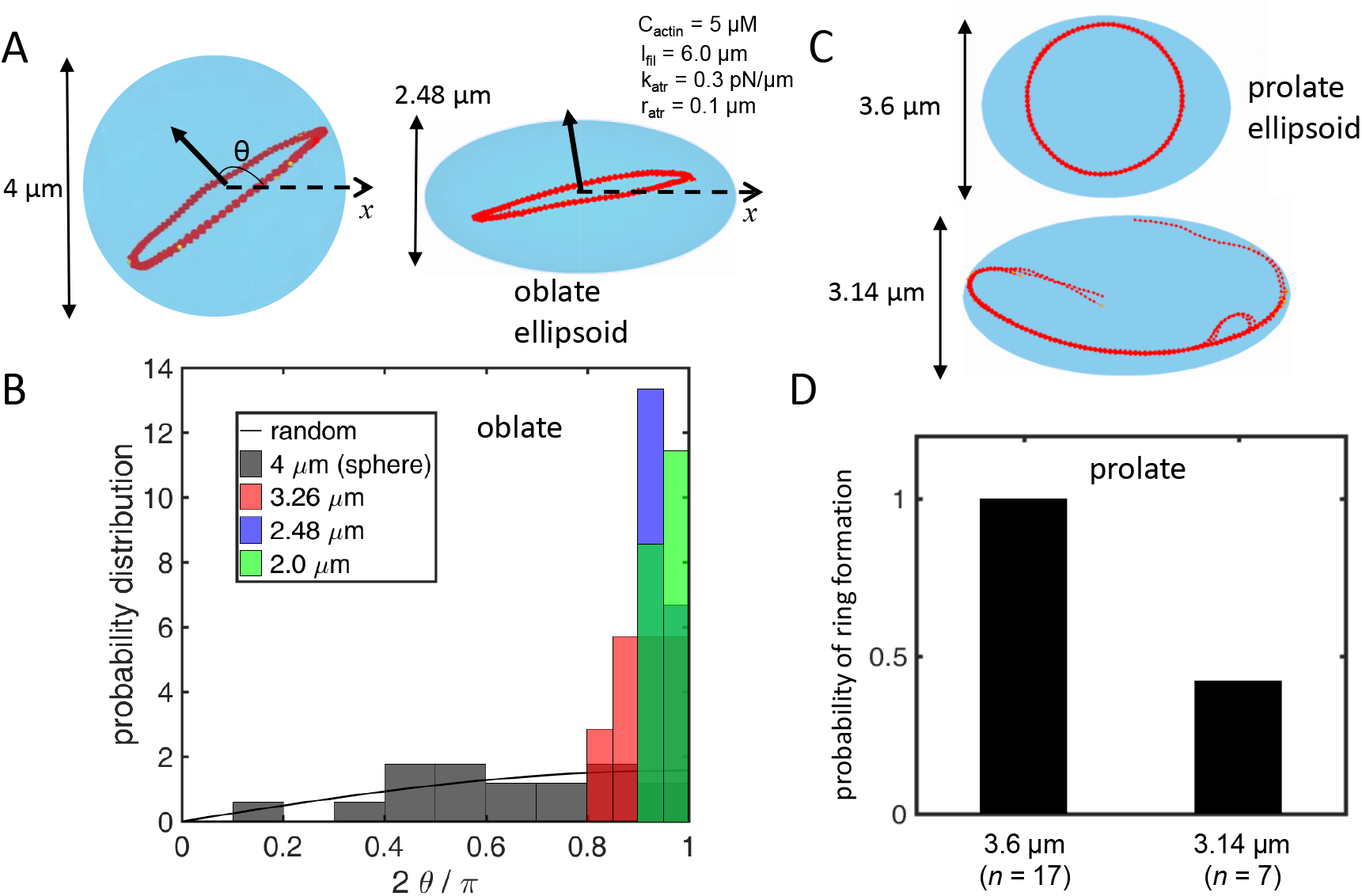
Ring formation and orientation in spheres and ellipsoids. Results of simulations (near 1000 *s*) for *C*_*actin*_ = 5 *µM*, all at the same volume, with 45 filaments reaching length *l*_*fil*_ = 6 *µm* for PS1 (*k*_*atr*_ = 0.3 *pN/µm, r_atr_* = 0.1 *µm*). (A) Snapshots of simulations. Left: sphere with *R*_*conf*_ = 2 *µm*. Right: oblate ellipsoid with same volume as sphere, semi-major axis 2.54 *µm* and semi-minor axis 1.24 *µm*. (B) Normalized distribution of angle between the average binormal vector shown in panel A and the *x* axis, for sphere (black, 17 runs) and ellipsoids (color, 7 runs each). Black line shows normalized sin function and inset labels indicate length of minor axis. (C) Snapshots of simulation for prolate ellipsoids with same volume as sphere in panel A. Top: semi-major axis 2.45 *µm* and semi-minor axis 1.8 *µm*. Bottom: semi-major axis 3.2 *µm* and semi-minor axis 1.57 *µm*. (D) Probability of ring formation for the two prolate ellipsoid geometries in panel C. A ring formed in all simulations with spheres and oblate ellipsoids in panels A and B.

We measured the angle *θ* between the vector perpendicular to the ring plane (determined by the average binormal vector over filament beads) and the fixed *x* axis that is along the long axis of the ellipsoid (Fig. 10A). For a random orientation distribution that is uniformly distributed along solid angle *d*Ω = sin *θ dθ dφ*, where *φ* is azimuthal angle, the distribution of *θ* would follow a sin curve. This is indeed close to our results (17 runs) for a sphere, which has no preferred orientation. By contrast, the orientation of rings forming in oblate ellipsoids becomes narrowly peaked at *θ* = *π/*2 with increased flattening. These results indicates a preference for the plane of the ring to align along the flat plane of the oblate ellipsoid such that bending energy cost is minimized at the expense of less contacts.

Simulations were also performed in prolate ellipsoids (Fig. 10 C,D). This geometry resembles the cylindrical shape of fission yeast. Like the oblate case, small deformations with respect to a spherical shape resulted in rings oriented such that the the major axis is along the ring plane (Fig. 10 C top, D). Larger deformations by contrast frequently resulted in an open ring/bundle that aligned along the major axis (Fig. 10 C bottom, D). The length of the major axis in the latter case was the largest among all ellipsoid simulations and comparable to the filament length.

#### 3.4.4 Attraction to confining boundary

So far we assumed that actin filament interactions with the confining boundary corresponded to a repulsive hard wall. However, depletion interactions by methylcellulose cause a short range attraction to the confining surface [Fisher and Kuo, 2008, Kang et al., 2016]. This may promote ring formation by opposing collapse of actin filaments toward the center of the confining sphere, as was shown to occur in Figs. 4-6 for the higher values of *k*_*atr*_ and *r*_*atr*_. To test the effect of surface attraction in these configurations, we introduced a short-range attraction force to the surface that still allows sliding along the boundary. We found that surface attraction can indeed oppose the collapse and promote ring formation (Fig. 11, S7). We suggest that this may be why increasing confining radius and concentration in Figs. 8C and 9C (that had no surface attraction) impacted ring formation more severely compared to experimental observations.

**Figure 11:**
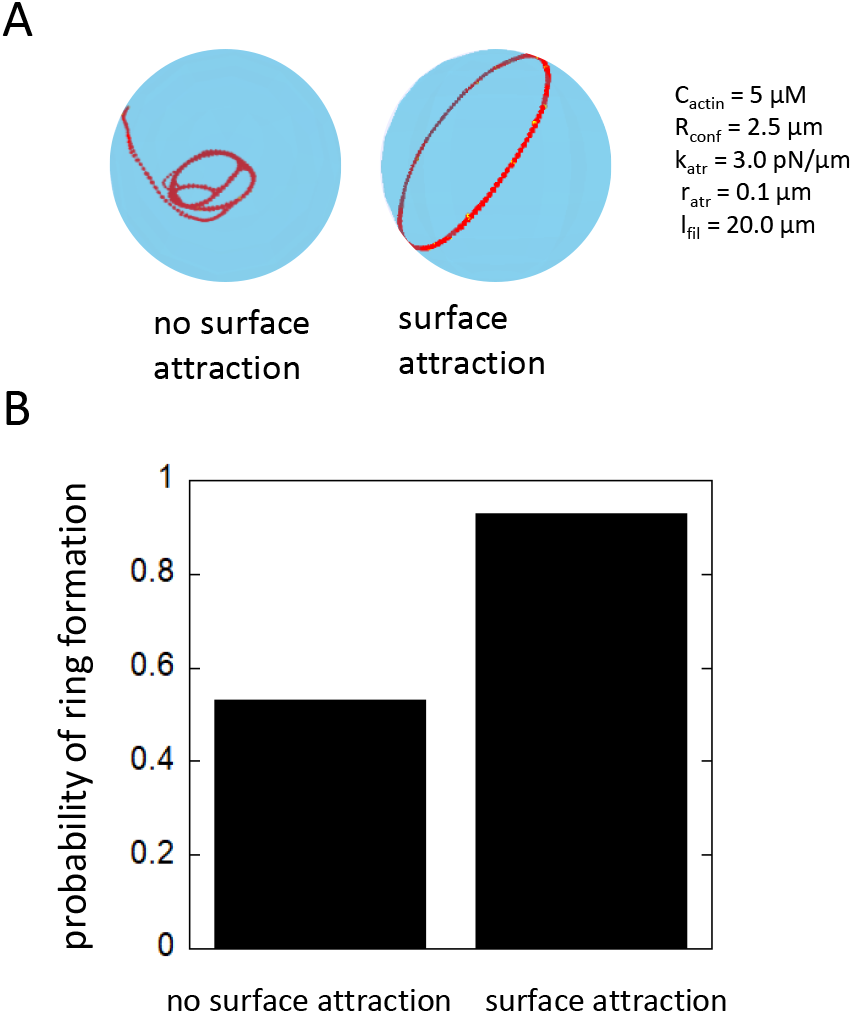
Simulations with surface attraction enhancing ring formation. (A) Snapshots of simulation at 1500 *s* for *R*_*conf*_ = 2.5 *µm* and *C*_*actin*_ = 5 *µM* with filaments reaching *l*_*fil*_ = 20 *µm*, *k*_*atr*_ = 3.0*pN/µm*, and *r*_*atr*_ = 0.1*µm* in the absence and presence of surface attraction. (B) Probability of ring or ring-like structure formation with and without surface attraction, out of 15 runs for each case. The difference is statistically significant (*p* = 0.035, Fisher’s exact test). Snaphsots for all runs shown in Fig. S7

## 4 Discussion

In this work, we performed simulations to study some of the main features of actin filament organization within confined spaces. We used a system with minimal elements that included the semi-flexible nature of actin filaments, their polymerization and diffusion, the effect of confinement, concentration, interactions between filaments and with the confining boundary. These Brownian dynamics simulations revealed that confinement can promote ring formation, however this occurs only within a optimal range of parameters values. Other structures such as bundled, curved open bundles, irregular loops, and small rings were also observed. In particular these structures depended on the kinetics of their assembly and on the ability of actin filaments to slide along filament bundles. Indeed, polymerization was found to be important for ring assembly in vitro [Miyazaki et al., 2015] while assembly kinetics also determine the structure of bulk actin networks [Falzone et al., 2012].

We found that generally, filaments of length much shorter than the size of confinement cannot form a ring in the absence of attraction to the surface. When filament length becomes comparable to, or longer than the size of confinement, the chance of ring formation increases. However, in this case the filament length also starts to become comparable to the actin persistence length and disorganized metastable arrangements (loops, collapsed, etc) can also occur. Prior models have described the formation of looped metastable minima structures for single self-attractive filaments [Ou and Muthukumar, 2005, Lappala and Terentjev, 2013]. Here we show how confinement leads to a shift from a kinetically-determined bulk network to rings and irregular loop structures. We note that such loop shapes might not form in the case of confined microtubules [Pinot et al., 2009], which are stiffer biopolymers. Crosslinked confined microtubule rings have been studied computationally in the context of regulation of blood cell shape and mechanics [Dmitrieff et al., 2017].

Our work further highlights how both geometry and mechanics as well as signaling and spatial patterning in the cell may need to coordinate for efficient actin organization For example, actomyosin ring assembly in fission yeast depends on cell geometry as well as correct placement and time of actin filaments near the plasma membrane [Bidone et al., 2014, Tang et al., 2015, Zhang et al., 2016]. Steps in this process involve the nucleation of actin filaments by formin proteins, compaction of actin filaments through myosin pulling and crosslinking by *α*-actinin and fimbrin proteins [Laporte et al., 2012].

Lim et al. [Lim et al., 2018] studied fission yeast spheroplasts that become spherical after degradation of their cell wall. Rings formed along the path of least curvature, in both compressed and uncompressed spheroplasts, even in cells that lacked certain spatial cues by proteins mid1 and tea1 for medial cytokinetic ring positioning [Lim et al., 2018]. Spheroplasts treated with actin-disrupting compound swindholide-A formed non-equoatorial actin rings [Lim et al., 2018]. Thus, these experiments further show a role for confining shape and filament length in ring formation, consistent with the results of this work and prior in vitro experiments. Further modeling and experimental work is needed to resolve the precise mechanism of ring formation in spheroplasts. In addition to cell geometry, we anticipate a role for formin actin filament nucleators, some of which assemble into membrane nodes in cells that lack mid1 [Saha and Pollard, 2012], recruitment and force generation of myosin recruited by actin to the membrane [Saha and Pollard, 2012], and actin filament turnover that may realize a search and capture process that depends on filament length [Vavylonis et al., 2008].

An advantage of our computational approach is that we can account for experimental values of confinement size, polymerization time, and actin concentrations. The attractive or crosslink interactions were treated in an approximate generic way, which however allowed us to explore a range of possible phenomena for both depletion and crosslink mediated interactions. We mainly considered the boundary as being a hard wall, even though depletion interactions would also induce attraction to the boundary and influence the resulting morphology (Fig. 11).

Fascin establishes stiff bundles whose thickness is limited by helical twist [Claessens et al., 2008]. Fascin-actin bundles can deform the soft confining boundary of liposomes [Honda et al., 1999, Tsai and Koenderink, 2015]. Tsai and Koenderink [Tsai and Koenderink, 2015] studied 10 *µM* actin in liposomes of 2 − 16 *µm* radius and 0.02 to 0.5 fascin-to-actin molar ratio. In non-deformed liposomes produced by increasing membrane rigidity, fascin-actin bundles organized into multiple or packed cortical rings. We speculate these morphologies are of similar origin to the irregular loop structures observed in Fig. 6, which form in a packed arrangement as a result of the increased bundle rigidity of fascin-actin bundles compared to the bundles in our simulations.

Another effect that may become important for densely aggregated structures, such as those in Fig. 6E, is filament breaking, which could result in compaction of such structures into short thick bundles. Finally, our free-draining approximation of filament diffusion results in the slowing-down of bundle diffusion compared to single filaments. Thus, the coalescence of bundles in Fig. 4 might occur faster than in simulations.

The discrete simulations performed in this work were useful to map out a complex interplay among mechanics, confinement, crosslink interactions that result in changes of macroscopic morphology. Future work that captures additional characteristics of each individual crosslinker and motor protein in a coarse-grained manner, together with related computational approaches [Nedelec and Foethke, 2007, Mak et al., 2016, Ma and Berro, 2018], can improve our understanding of how the cells assemble and construct functional cytoskeletal structures.

## Acknowledgments

We thank Gijsje Koenderink and Feng Tsai for discussions and feedback that motivated this work. We also thank Haosu Tang for help with simulations. This work was supported by National Institutes of Health Grant R01GM114201. Use of the high-performance computing capabilities of the Extreme Science and Engineering Discovery Environment (XSEDE), which is supported by the National Science Foundation, project no. TG-MCB180021 is also gratefully acknowledged.

## Data availability

The data and code that support the findings of this work are available from the corresponding author upon reasonable request.

