## Supplementary Material for "Organization of Associating or Crosslinked Actin Filaments in Confinement"

### Supplementary Movies and Figures: Organization of Associating or Crosslinked Actin Filaments in Confinement

#### Movies

Movie S1. Movie of simulation showing formation of irregular aggregate for $C_{actin} = 5\,\mu M$, confining radius $R_{conf} = 2.5\,\mu m$, final filament length $l_{fil} = 1.2\,\mu m$ and interaction parameters $k_{atr} = 3\,pN/nm$, $r_{atr} = 0.06\,\mu m$. The time between frames is 10 $s$ and total time 1000 $s$. The yellow beads show barbed (polymerizing) end of filament.

Movie S2. Movie of simulation showing ring formation for $C_{actin} = 5\,\mu M$, confining radius $R_{conf} = 2.5\,\mu m$, final filament length $l_{fil} = 3.8\,\mu m$ and interaction parameters $k_{atr} = 2\,pN/nm$, $r_{atr} = 0.08\,\mu m$. The time between frames is 10 $s$ and total time 500 $s$.

Movie S3. Movie of simulation showing open bundle formation for $C_{actin} = 5\,\mu M$, confining radius $R_{conf} = 2.5\,\mu m$, final filament length $l_{fil} = 3.8\,\mu m$ and interaction parameters $k_{atr} = 0.3\,pN/nm$, $r_{atr} = 0.1\,\mu m$. The time between frames is 10 $s$ and total time 1000 $s$.

Movie S4. Movie of simulation showing formation or irregular/collapsed loop for $C_{actin} = 5\,\mu M$, confining radius $R_{conf} = 2.5\,\mu m$, final filament length $l_{fil} = 20\,\mu m$ and interaction parameters $k_{atr} = 3\,pN/nm$, $r_{atr} = 0.1\,\mu m$. The time between frames is 10 $s$ and total time 500 $s$.

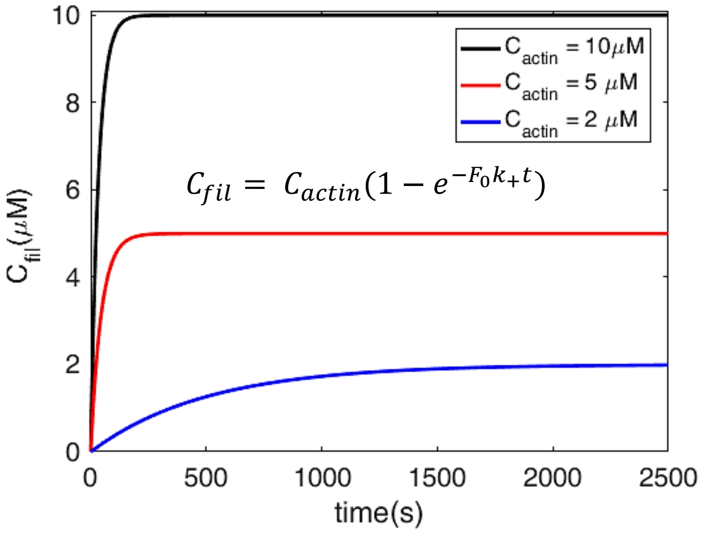


Figure S1: Plot of assumed dependence of actin filamentous concentration as a function of time. Here $k_+$ is the actin filament barbed end polymerization rate constant (see Table 1 of main text). The different curves show different assumptions on the initial concentration of actin filament seeds, $F_0$. In this figure, the filament seed concentration for total actin concentration $C_{actin} = 2\,\mu M$ (blue) was set to $F_0 = 3\,nM$; for $C_{actin} = 10\,\mu M$ (black) it was set to $F_0 = 0.2\,nM$. Using these values of $F_0$, the resulting curves approximately capture the measured polymerization curves in the experiments by Miyazaki et al. (2015). Since this simple model does not include a nucleation stage, the curves do not show an initial lag that would be more noticeable for the $2\,\mu M$ case [Miyazaki et al., 2015]. In this paper we used an intermediate concentration $C_{actin} = 5\,\mu M$ (red). For this intermediate case we set $F_0 = 2.2\,nM$, giving a polymerization curve in between the black and blue curves and which also gives a final actin filament length $l_{fil} = 6\,\mu m$, close to $4.6\,\mu m$ measured in droplets by Miyazaki et al. at 10 $\mu M$ concentration.

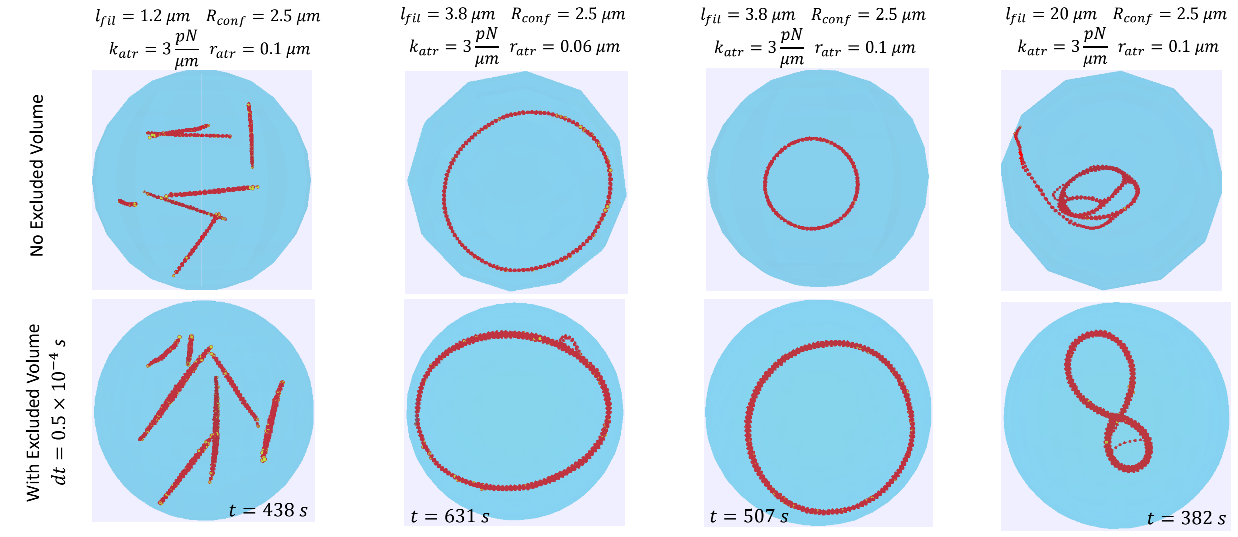


Figure S2: Snapshots of simulations without (top row) and with (bottom row) excluded volume interactions. All simulations have $C_{actin} = 5\,\mu M$, other parameters shown at the top of each column. The number of filaments in each column was 420, 140, 140 and 26, respectively. Snapshots without excluded volume shown at $1500\,s$. Snapshots with excluded volume were performed with a $dt$ that was three times smaller and shown at indicated times.

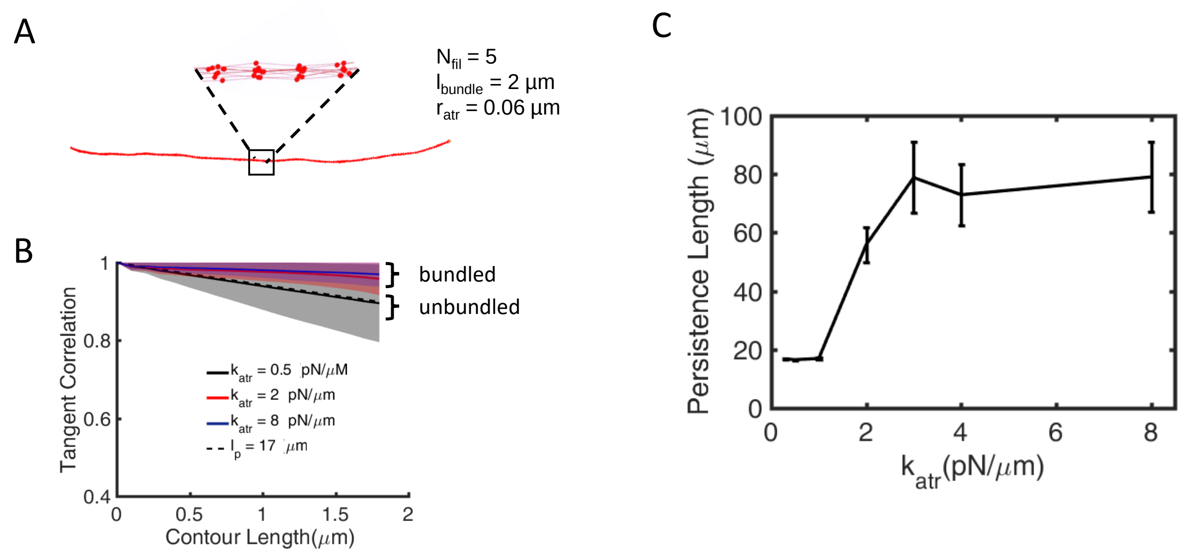


Figure S3: Persistence length of simulated bundles. (A) Snapshot of a bundle of five filaments of length $2\,\mu m$ each, interacting with each other via an attractive potential with $r_{atr} = 0.06\mu m$ and $k_{atr} = 8.0 pN/\mu m$. Enlarged region shows the locations of the actin filament beads in the bundle. The bundle is generated by polymerizing filaments along parallel directions and close enough to each other. (B) Tangent correlation for a filament in a bundle of of five filaments of $2\,\mu m$ each, measured by sampling over each filament for 2000 s for $r_{atr} = 0.06\mu m$ and various $k_{atr}$. The error bar (shaded area) shows standard deviation. For small values of $k_{atr}$ the filaments unbundle. (C) Persistence length of bundle of five filaments of $2\,\mu m$ over for 2000 $s$ after polymerization, as a function of $k_{atr}$ for $r_{atr} = 0.06\mu m$. The persistence length is found by an exponential fit to the curves of panel B as in Tang et al. (2014). Error bar shows standard error of fit.

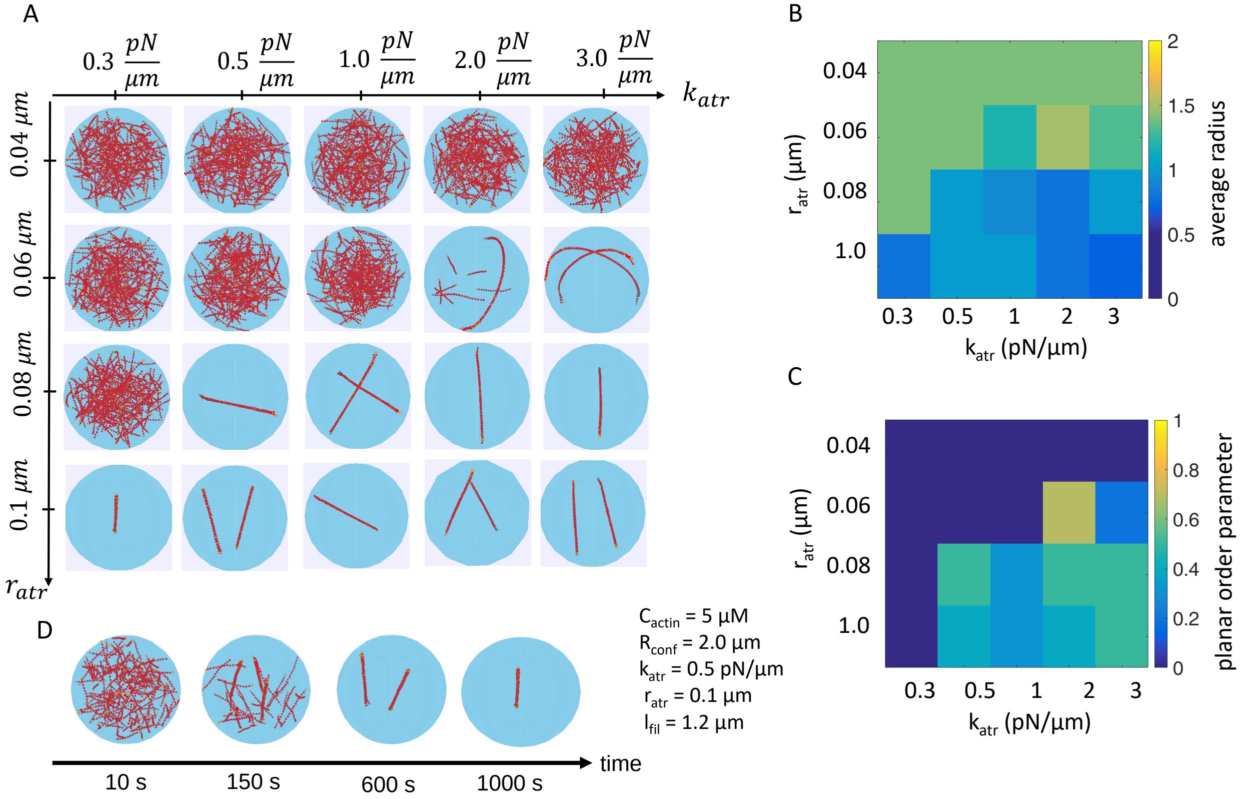


Figure S4: Actin network morphology for filaments that elongate to lengths shorter than the diameter of the confinement. Similar to Fig. 4, for a smaller droplet of $R_{conf} = 2\,\mu m$. (A) Snapshots of simulation at $1500\,s$ for $C_{actin} = 5\,\mu M$, 180 filaments, and final filament length $l_{fil} = 1.4\,\mu m$ for four values of $r_{atr}$ and four values of $k_{atr}$ as indicated. (B) Average filament bead radius, sampled from 1000 to $1500\,s$ every $10\,s$, as a function of $r_{atr}$ and $k_{atr}$, one run. (C) Average planar order parameter, sampled as in panel B. (D) Snapshots of the case with $k_{atr} = 0.5pN/\mu m$ and $r_{atr} = 0.1\mu m$ over time.

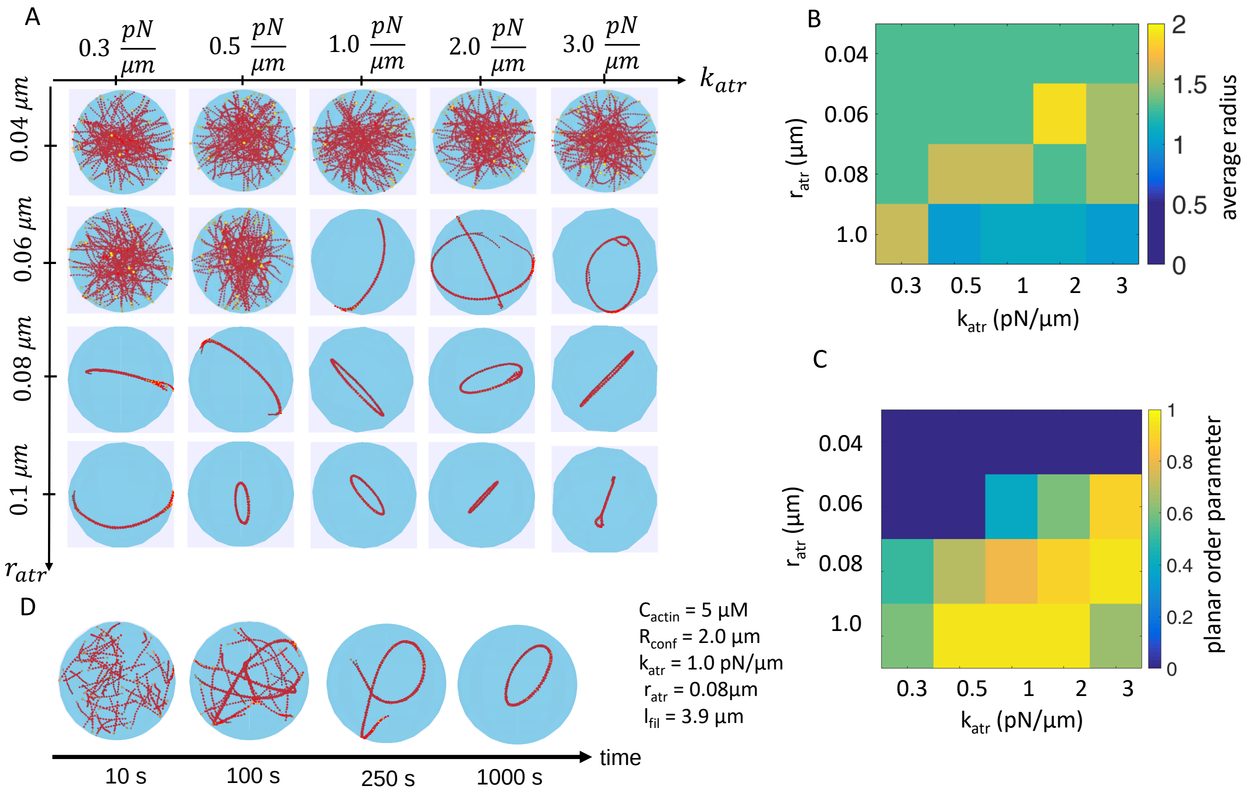


Figure S5: Actin network morphology for filaments that elongate to lengths comparable to the diameter of the confinement. Similar to Fig. 5, for a smaller droplet of $R_{conf} = 2\,\mu m$. (A) Snapshots of simulation at $1500\,s$ for $C_{actin} = 5\,\mu M$, and 69 filaments reaching $l_{fil} = 3.8\,\mu m$. (B) Average radius, sampled from 1000 to 1500 $s$ every $10\,s$, one run. (C) As panel B, for planar order parameter. (D) Snapshots of the case $k_{atr} = 1\,pN/\mu m$ and $r_{atr} = 0.08\,\mu m$.

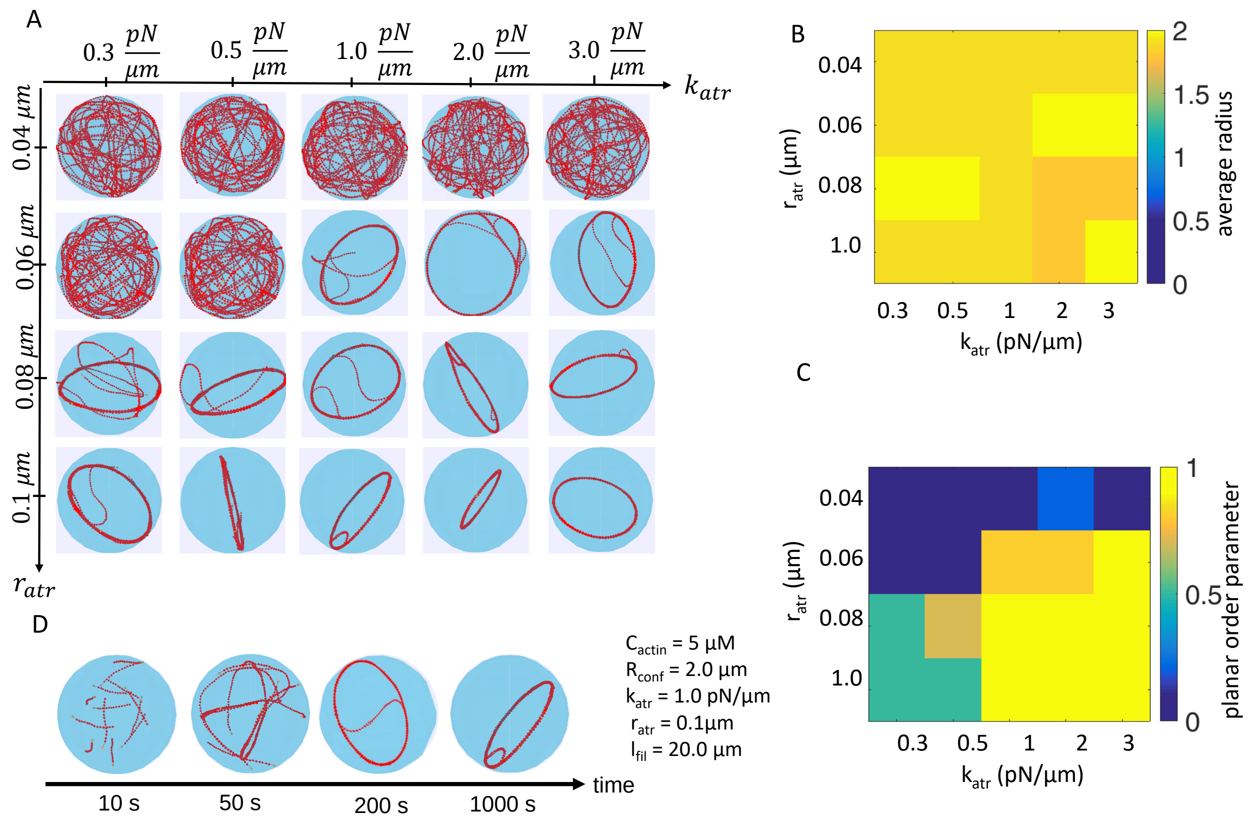


Figure S6: Actin network morphology for filaments that elongate to lengths longer than the diameter of the confinement. Similar to Fig. 6, for a smaller droplet of $R_{conf} = 2\,\mu m$. (A) Snapshots of simulation at $1000\,s$ for $C_{actin} = 5\,\mu M$, and 20 filaments reaching $l_{fil} = 20\,\mu m$. (B) Average radius, sampled from 1000 to 1500 $s$ every $10\,s$, one run. (C) As panel B, for planar order parameter. (D) Snapshots of the case $k_{atr} = 1\,pN/\mu m$ and $r_{atr} = 0.1\,\mu m$.

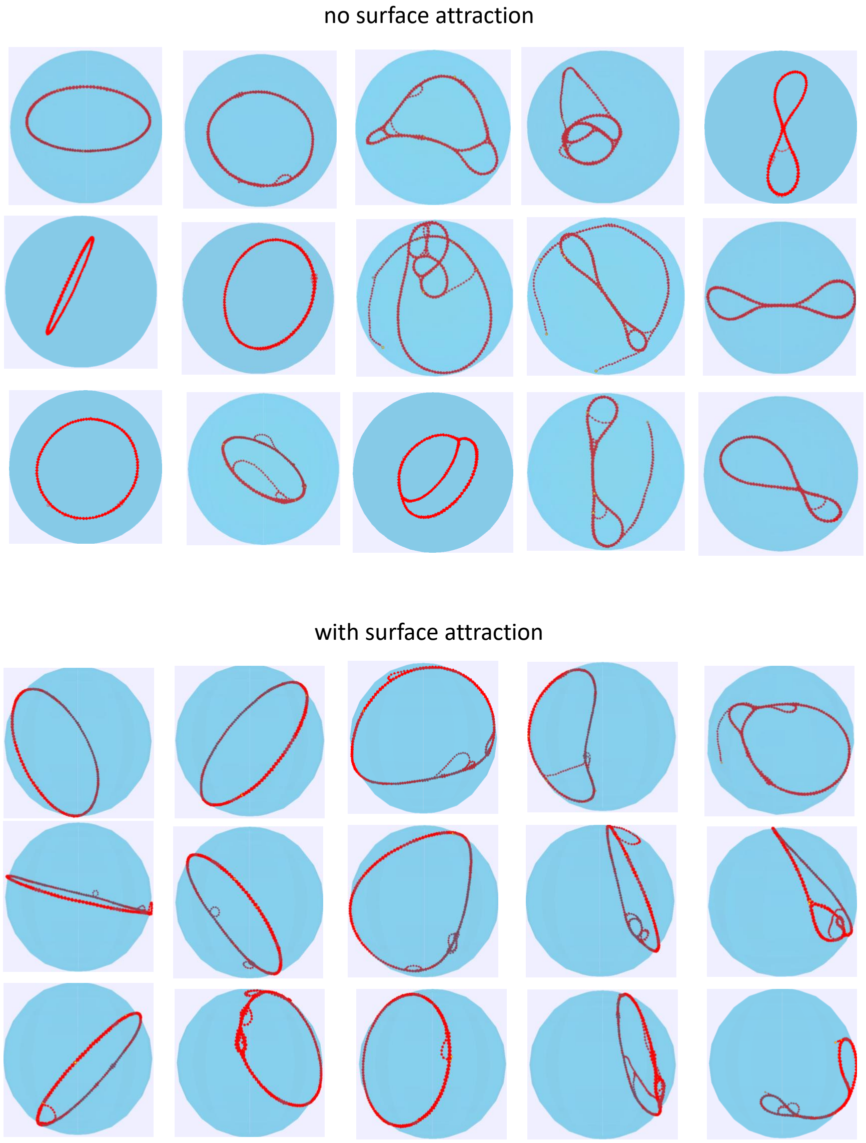


Figure S7: Snapshots of simulation results (for times above 750 s) with and without surface attraction. Thes simulations correspond to Fig. 11 where $R_{conf} = 2.5\,\mu m$, $C_{actin} = 5\,\mu M$, filaments reaching $l_{fil} = 20\,\mu m$, $k_{atr} = 3.0 pN/\mu m$, and $r_{atr} = 0.1 \mu m$.

| Parameter | Fig. 1D | Fig. 2A | Fig. 2C | Fig. 3 |
|---|---|---|---|---|
| $R_{conf}$ ($\mu m$) | 2.5 | 2.5 | 2.5 | no confinement |
| $C_{actin}$ ($\mu M$) | 5 | varied | 11 | 5 |
| $N_{fil}$ | 140 | 60 | 60 | 256 |
| $l_{fil}$ ($\mu m$) | 3.8 | varied | 20 | 3.8 |
| $k_{atr}$ ($pN/\mu m$) | 2 | no attraction | no/3 | varied |
| $r_{atr}$ ($\mu m$) | 0.08 | no attraction | no/0.06 | 0.06 |
| surface attraction | no | no | no | N/A |

| Parameter | Fig. 4 | Fig. 5 | Fig. 6 | Fig. 7 |
|---|---|---|---|---|
| $R_{conf}$ ($\mu m$) | 2.5 | 2.5 | 2.5 | summary of |
| $C_{actin}$ ($\mu M$) | 5 | 5 | 5 | Figs. 4-6 and |
| $N_{fil}$ | 420 | 140 | 26 | Figs. S4-S6 |
| $l_{fil}$ ($\mu m$) | 1.2 | 3.8 | 20 | |
| $k_{atr}$ ($pN/\mu m$) | varied | varied | varied | |
| $r_{atr}$ ($\mu m$) | varied | varied | varied | |
| surface attraction | no | no | no | |

| Parameter | Fig. 8 | Fig. 9 | Fig. 10 | Fig. 11 |
|---|---|---|---|---|
| $R_{conf}$ ($\mu m$) | varied | 2.5 | 2/shape varied | 2.5 |
| $C_{actin}$ ($\mu M$) | 5 | varied | 5 | 5 |
| $N_{fil}$ | varied | varied | 45 | 26 |
| $l_{fil}$ ($\mu m$) | 6 | 6 | 6 | 20 |
| $k_{atr}$ ($pN/\mu m$) | 0.3/3 | 0.3/3 | 0.3 | 0.3 |
| $r_{atr}$ ($\mu m$) | 0.1/0.08 | 0.1/0.08 | 0.1 | 0.1 |
| surface attraction | no | no | no | yes/no |

| Parameter | Fig. S4 | Fig. S5 | Fig. S6 | Fig. S7 |
|---|---|---|---|---|
| $R_{conf}$ ($\mu m$) | 2 | 2 | 2 | 2.5 |
| $C_{actin}$ ($\mu M$) | 5 | 5 | 5 | 5 |
| $N_{fil}$ | 180 | 69 | 20 | 26 |
| $l_{fil}$ ($\mu m$) | 1.4 | 3.8 | 20 | 20 |
| $k_{atr}$ ($pN/\mu m$) | varied | varied | varied | 0.3 |
| $r_{atr}$ ($\mu m$) | varied | varied | varied | 0.1 |
| surface attraction | no | no | no | yes/no |

Table S1: Summary of Parameter Values Used in Figures. Parameters not listed were taken from Table 1.
